# Rhizobacterial Biosensors Spatially Map Natural and Engineered Sucrose Exudation

**DOI:** 10.64898/2026.04.15.718812

**Authors:** Christopher M. Dundas, Gretchen A. Brinkman, Taylor Clarke, Madison Payne, José Alama Ureta, Ivy Velasco, Jason G. Wallace, José R. Dinneny

## Abstract

Root exudation mediates the delivery of plant primary and secondary metabolites into soil, where they regulate plant–microbe interactions and terrestrial carbon cycling. Conventional exudate analyses quantify total root-released carbon yet obscure the spatial origin and rhizosphere influence of individual compounds. Here, we develop a rhizobacterial biosensor platform, named Suc-MAPP, to map local exudate profiles along the surface of colonized root tissues. Focusing on sucrose, we engineered sfGFP-based, sucrose-responsive gene circuits in *Pseudomonas putida* KT2440 for live imaging of exudate concentrations in the micromolar range. These biosensors reveal spatially structured sucrose exudation patterns across eudicots and monocots and implicate photoassimilated source–sink dynamics as a major determinant. We further apply this platform to phenotype exudation modulated by synthetic gene circuitry in *Arabidopsis thaliana*, identifying genetic design rules for graded sucrose release and quantifying how engineered export sculpts rhizosphere assembly of a defined bacterial community. Together, these results establish programmable rhizobacterial biosensors as tools to spatially resolve plant–environment carbon exchange *in situ* and provide a framework for extending this approach to diverse exudate targets.

## Main

Plant roots direct 20–40% of their photosynthates into the surrounding soil (rhizosphere) as a diverse array of organic compounds, termed root exudates^1,2^. These metabolites drive plant nutrient acquisition^3^, soil organic matter formation^4^, and serve as carbon sources for root-associated microbiota^5^. Increasing evidence highlights the presence of spatially-distinct exudate distributions that shape rhizosphere community assembly^6,7^. Because these patterns govern phytopathogen and plant-growth-promoting microbial activities, precise localization of exudates through natural and engineered variation has significant potential to optimize agriculture, environmental biotechnology, and soil carbon sequestration^8^. However, the spatiotemporal dynamics and physiological controllers for most root-released compounds remain unclear. Establishing a tissue-scale understanding of exudation will be critical to link rhizosphere molecular distributions to plant-recruitment of microbes under environmental stress (e.g., the “cry-for-help” hypothesis^9^) and enable the design of targeted host-microbe cross-feeding for improved biofertilizer efficacy and biocontainment^10^.

This knowledge gap largely stems from the standard protocols used for root exudate analysis: sampling bulk exudates from roots submerged in well-mixed hydroponic systems. These approaches aggregate total root-released carbon for metabolomic profiling, and have revealed the chemical diversity of exudates and their broad influence on microbiome assembly^11,12^. However, since these methods lack spatial resolution, they cannot resolve locations of metabolite release^13^, map exudate distributions across root tissues and developmental zones^14^, nor capture fine-scale rhizosphere ecologies such as localized carbon competition^15,16^. This contrasts with recent advances in plant single-cell and spatial analysis approaches that resolve transcriptome and proteome maps across cell-types and root tissues^17–19^. To enable comparable resolution for studying and engineering rhizosphere carbon dynamics, new methods are needed that can quantify extracellular plant metabolites *in situ*.

Genetically encoded sensors enable sensitive, specific, and quantitative detection of environmental analytes^20^, and represent a promising yet underutilized tool for spatial mapping of root exudates^21^. Whole-cell microbial biosensors (i.e., systems with intact, live cells) are attractive for plant applications due to their programmable ligand-responsive transcriptional elements^22,23^ and ability to detect exudation across colonized tissues^24–27^. However, previously developed systems rely on promoter–reporter fusions lacking exudate specificity^26^ or on microbial chassis with restricted colonization domains (e.g., root-nodulating rhizobia^27^), thus limiting spatial resolution of biosensing. The expanding synthetic biology toolkits for both rhizobacteria and plants^28^ creates an opportunity to optimize these technologies and leverage transkingdom engineering approaches to both map and manipulate rhizosphere carbon in a spatially resolved manner^27,29^.

We hypothesized that microbial-based whole-cell sucrose biosensing in the rhizosphere would be particularly valuable due to the sugar’s high abundance in root exudates^11,30^ and strong connection to photosynthesis and source–sink carbon allocation^31^. Prior studies observed pronounced differences in bulk-collected sucrose exudates across different developmental timescales^11,30^, suggesting that biosensors might resolve spatially localized exudation patterns across root developmental zones. While a whole-cell sucrose biosensor relying on bacterial repressor ScrR was previously reported^24,32^, other studies suggest this protein does not detect sucrose but instead responds to the sucrose catabolism byproduct and abundant rhizosphere exudate, fructose^33–35^. Despite advances in creating rhizobacterial biosensors for other sugars and rhizosphere metabolites^36^, and the creation of FRET-based sucrose sensing proteins^37^, sucrose-specific rhizobacterial biosensors remain underdeveloped.

Here, we engineered a fluorescent rhizobacterial biosensor platform termed Suc-MAPP (Sucrose-Metabolite Abundance Pattern Profiler) to map root sucrose exudation, interrogate species variation, and test synthetic sucrose exudation circuits in *A. thaliana*, linking these changes to microbiome assembly (Figure 1a). The modularity of our bacterial and plant genetic circuits, combined with a scalable fluorescence imaging workflow, establishes a generalizable strategy for quantifying and tuning root exudation and its impacts on plant physiology and rhizosphere interactions.

**Figure 1.**
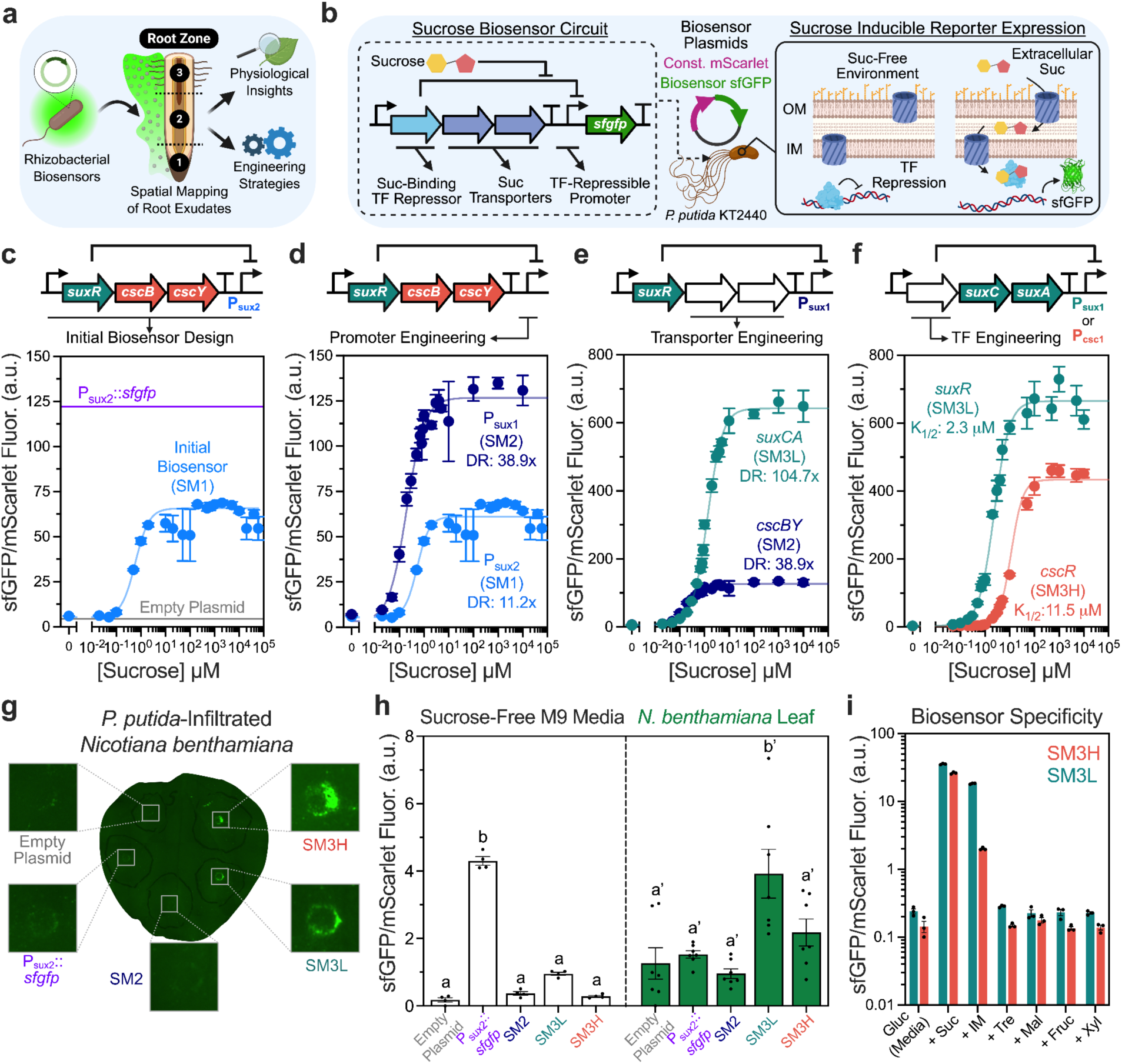
Development of Sucrose Biosensors. **a,** A schematic of rhizobacterial biosensors mapping root zone exudation, enabling study of plant physiology and exudate engineering. **b,** A schematic of sucrose biosensor circuit design, deployed in *P. putida* that relies on sucrose uptake and transcriptional derepression to produce sfGFP. **c–f** Response functions of Suc-MAPP variants (selected curves replotted for comparison). **c,** Initial biosensor variant, Suc-MAPP-1 (SM1) **d,** Promoter engineering, generating Suc-MAPP-2 (SM2). **e,** Transporter engineering, generating Suc-MAPP-3L (SM3L) **f,** Repressor transcription factor engineering, generating Suc-MAPP-3H (SM3H). All response function data represent mean ± SEM of n = 3. **g,** Example *Nicotiana benthamiana* leaf infiltrated with SM3H (orange), SM3L (teal), SM2 (dark blue), P_sux2_::*sfgfp* (purple) *P. putida* biosensors, and an empty plasmid control. Rings indicate the site of infiltration. **h,** Quantification of *P. putida* biosensor signal (sfGFP/mScarlet fluorescence) in liquid sucrose-free M9 media (white bars) and *N. benthamiana* leaf discs (green bars). Significance groups were determined by one-way ANOVA with post-hoc Tukey’s, P < 0.05. n = 4 (sucrose-free M9), n = 7 (leaf infiltration disc) biological replicates. Data represent mean ± SEM. **i,** Quantification of SM3H (orange) and SM3L (teal) biosensor response to a panel of di- and monosaccharides. Data represent mean ± SEM, n = 3 biological replicates. Gluc = glucose, Suc = sucrose, IM = isomaltulose, Tre = trehalose, Mal = maltose, Fruc = fructose, Xyl = xylose.

## Results

### Development of Bacterial Sucrose Biosensor Circuits

We engineered rhizobacterium *Pseudomonas putida* KT2440 as a tunable whole-cell sucrose biosensor, leveraging its well-annotated genetics^38^, synthetic biology toolkits^39,40^, and capacity for root-colonization^41,42^. KT2440 lacks native sucrose uptake and catabolism^43^ (Figure S1), enabling rhizosphere sucrose detection without perturbing local concentrations through consumption. Biosensor circuits were built using LacI-family sucrose-response transcriptional repressors (SuxR or CscR), outer (CscY or SuxA) and inner (CscB or SuxC) membrane sucrose transporters, and a synthetic SuxR- or CscR-regulated promoter driving sfGFP expression. These were expressed on broad-host-range plasmids, along with a constitutive mScarlet reporter, enabling ratiometric sfGFP/mScarlet fluorescence to serve as quantitative measurements of local sucrose concentration^43,44^ (Figure 1b).

Systematic optimization of biosensor-operon promoter and ribosome binding site strengths, output promoter architecture, transporters, and repressor identity produced a series of *P. putida* biosensor variants with tunable sucrose detection (Figure 1c–f, Figure S2–S3). *In vitro* induction across a gradient of sucrose concentrations was well-described by a sigmoidal, Hill-type function, where dynamic range (DR) reflects fold-change between uninduced and induced states, and K_1/2_ represents the concentration required for half-maximal activation (i.e., sensor sensitivity) (Methods). Variants exhibited robust induction to micromolar sucrose above empty vector controls and spanned a broad range of responses: Suc-MAPP-1 (DR = 12.4x, K_1/2_ = 0.59 μM), Suc-MAPP-2 (DR = 38.9x, K_1/2_ = 0.20 μM), Suc-MAPP-3L (DR = 104.7x, K_1/2_ = 2.3 μM), and Suc-MAPP-3H (DR = 144.5x, K_1/2_ = 11.5 μM). Notably, Suc-MAPP-3L and Suc-MAPP-3H exhibited the highest dynamic ranges (>100x), with Suc-MAPP-3H shifting the detection window toward higher sucrose concentrations (Figure 1f). Profiling of these circuits in *Paraburkholderia graminis* confirmed cross-host portability with differing sensing thresholds (Figure S4), demonstrating flexible sucrose biosensor deployment across rhizobacteria in two different taxonomic classes (Gammaproteobacteria and Betaproteobacteria).

*In planta* biosensor functionality was confirmed using the *Nicotiana benthamiana* leaf apoplast, which supports *Pseudomonas* species colonization^45^ and contains millimolar sucrose (0.4–5.0 mM^46,47^). Leaf-infiltrated Suc-MAPP-3L and Suc-MAPP-3H showed an increase in ratiometric signal similar to sucrose-supplemented media (Figure 1g; Figure S5), whereas all other circuits were indistinguishable from empty plasmid controls in sucrose-free media (Figure 1h). Fluorescence microscopy showed that biosensor activation localized to infiltration sites (Figure 1g), consistent with tissue damage-induced increases in apoplastic sucrose^48^. Notably, Suc-MAPP-2 did not induce *in planta*, though it only differs from Suc-MAPP-3L in its sucrose transporter identities. This aligns with predicted sucrose transporter affinities, where CscBY (SucMAPP-2) exhibits millimolar sucrose affinities^43^ while SuxAC (Suc-MAPP-3L) is in the micromolar range^44^ (Figure 1e), further supporting plant-derived sucrose as the biosensor activation source. Indeed, Suc-MAPP-3L and Suc-MAPP-3H showed no induction with other plant sugars and only partial induction with isomaltulose (IM), a sucrose isomer rarely found in plant tissues^49^ (Figure 1i). Together, these results establish Suc-MAPP-3L and Suc-MAPP-3H as sucrose-specific, *in planta* biosensors.

### Sucrose Biosensors Define Spatially Localized Root Exudation Patterns

We deployed *P. putida* biosensors in the *A. thaliana* Col-0 rhizosphere to map sucrose exudation along the primary root. Axenic seedlings grown on solid MS media were root tip-inoculated with Suc-MAPP-3L, Suc-MAPP-3H, or control strains, and imaged *in situ* 5 days post inoculation (dpi) (Figure 2a). sfGFP and mScarlet fluorescence localized to the surface of root tissues, consistent with known *P. putida* colonization of *A. thaliana* Col-0^29^, where control constructs yielded consistently high (P_Sux2_::*sfgfp*) or low (Empty Vector) ratiometric reporter signals along the primary root. In contrast, Suc-MAPP-3L revealed a spatial gradient of biosensor activation, with a low fluorescence ratio at the root tip that monotonically increased toward mature tissues (Figures 2b–c). Suc-MAPP-3H remained uninduced across developmentally young root regions and activated only beyond ∼4 cm from the root tip (Figure 2c), consistent with its higher K_1/2_ activation threshold (Figure 1f). mScarlet fluorescence, a proxy for bacterial abundance, was consistent across root developmental zones and between biosensor variants (Figure 2c), indicating that biosensor-determined exudation patterns do not arise from differential colonization. Meta-analysis of multiple *P. putida* Suc-MAPP-3L–Col-0 experiments from two institutions using the same microscope model showed that, despite variation in absolute biosensor activation, the pattern of low root tip signal was relatively consistent (Figure S6). We also inoculated Col-0 with Suc-MAPP-3L in rhizobacterium *Paraburkholderia gramini*s which revealed similar biosensor patterns to *P. putida* (Figure S7). As biosensor activation by *P. putida* on sucrose-supplemented plant-free solid media (Figures S8) was achieved at similar concentrations to liquid experiments (Figure 1f), these collective results suggest that Suc-MAPP-3L reports low micromolar sucrose gradients (0.1–10 μM) in the Col-0 rhizosphere.

**Figure 2.**
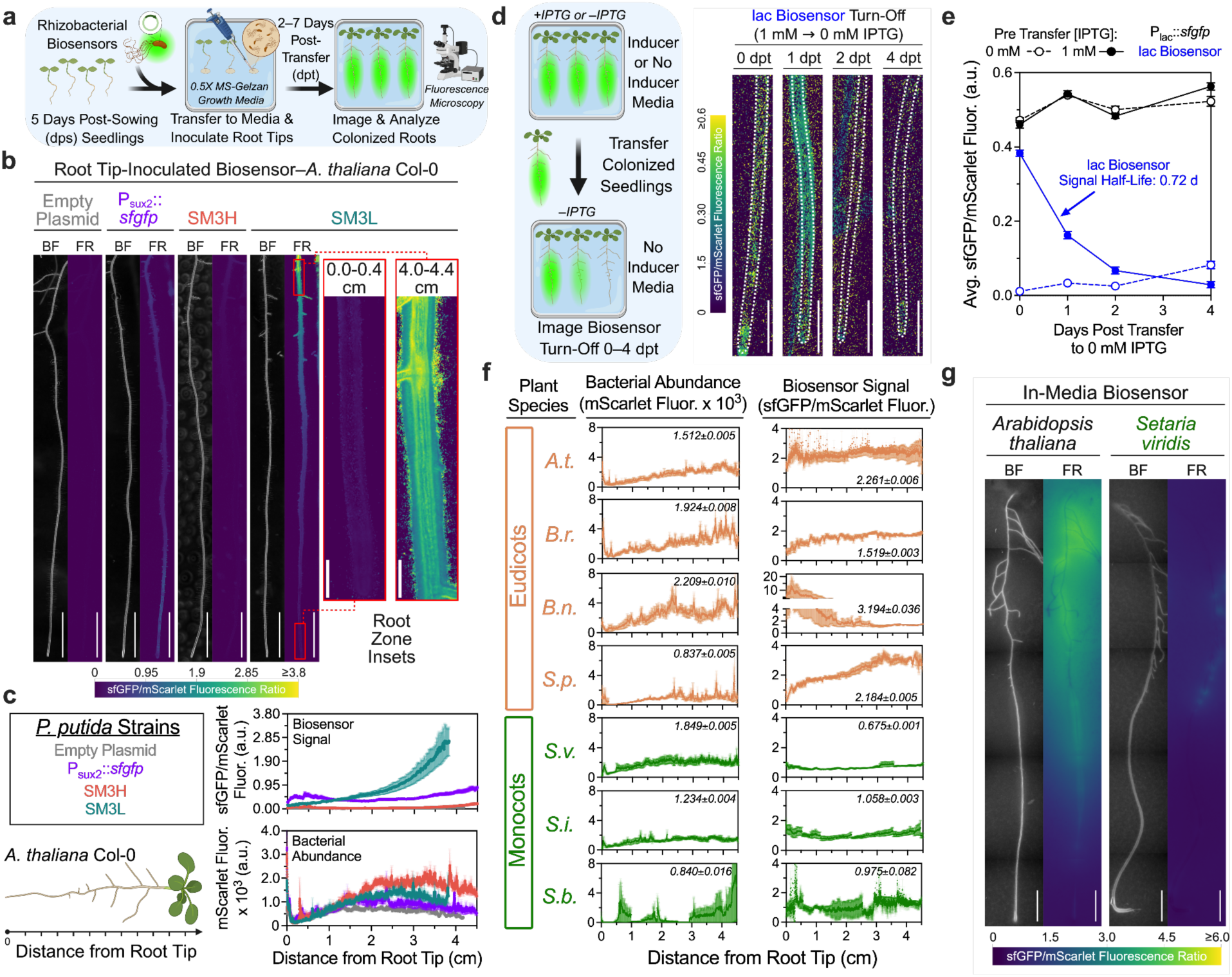
Defining Sucrose Exudation Patterns Across Roots. **a,** Experimental outline highlighting the primary root biosensor inoculation protocol. Seedlings 5 dps were inoculated with *P. putida* bacterial biosensors upon transfer to sucrose-free plant medium then imaged 2-7 days later. **b,c,** Sucrose signal output divergence across developed biosensor variants within living root systems. Representative images displaying sfGFP/mScarlet fluorescence along *A. thaliana* Col-0 primary roots inoculated with P_sux2_::*sfgfp* (purple), Suc-MAPP-3H (SM3H, orange), Suc-MAPP-3L (SM3L, teal), and an empty plasmid control. Insets highlight the SM3L signal measured at the root tip (0.0–0.4 cm) and maturation (4.0–4.4 cm) zones. Roots were imaged 5 dpi. Scale bar, 1 cm. **c,** Quantification of sfGFP/mScarlet fluorescence (mean ± SEM) and mScarlet fluorescence (mean ± SEM) along whole roots. **d,e,** Experimental outline of biosensor circuit turn-off assay. Seedlings inoculated with an IPTG-inducible variant of the Suc-MAPP-3L/HI were transferred from IPTG+/− medium to IPTG-medium to measure the timeline for signal turn-off. Images display sfGFP/mScarlet fluorescence along a root tip section (0–0.5 cm) of *A. thaliana* Col-0 inoculated with the *lac* biosensor imaged 0-4 dpt. Root traces from brightfield images are overlaid as dashed lines. Scale bar, 1000 µm. **e,** Quantification of average sfGFP/mScarlet fluorescence (mean ± SEM) along whole roots 0–4 dpt from IPTG-(open circles) or IPTG+ (closed circles) medium. Black lines represent the P_lac_::*sfgfp* positive control while blue lines represent the IPTG-inducible *lac* biosensor. **f,** Quantification of SM3L sfGFP/mScarlet fluorescence (mean ± SEM) and mScarlet fluorescence (mean ± SEM) along the primary roots of a multispecies plant panel including eudicots (orange) and monocots (green). Plants shown include *Arabidopsis thaliana* (*A.t.*), *Brassica rapa* (*B.r.*), *Brassica napus* (*B.n.*), *Solanum pennellii* (*S.p.*), *Setaria viridis* (*S.v.*), *Setaria italica* (*S.i.*), *Sorghum bicolor* (*S.b.*). Whole-root averages of sfGFP/mScarlet fluorescence ratio and mScarlet fluorescence (mean ± SEM) are indicated within each graph in italics. **g,** Example images of 10 day old *A. thaliana* Col-0 (black) and *S. viridis* (green) seedlings imaged 24 hours post transfer to SM3L-embedded solid medium. Scale bar, 0.5 cm.

We next probed the temporal dynamics of biosensor activation across root conditions. For roots inoculated with *P. putida* Suc-MAPP-3L at the same initial timepoint (5 days post sowing, dps), the biosensor showed distinct spatial patterns emerging between younger (2 dpi) and older (5–7 dpi) roots (Figure S9), where later timescales appear relatively comparable. To assess general features of biosensor signal decay in root-colonizing bacteria, we tested an orthogonally inducible *P. putida* construct responsive to IPTG (Figure S10), termed the *lac* Biosensor. *lac* Biosensor-colonized Col-0 seedlings were first grown in the presence of sensor-inducing IPTG, then transferred to inducer-free media where timelapse imaging confirmed exponential sensor turn-off (t_1/2_ = 0.72 d) (Figures 2d–2e). Extrapolating results to Suc-MAPP-3L, biosensor activation reflects dynamic responses to rhizosphere sucrose within an approximate one-day timescale, rather than irreversible sensor activation.

Given the broad colonization range of *P. putida*^29,41,42^, we hypothesized that sucrose biosensors could compare root exudation across plant species. Using root tip-inoculation with *P. putida* Suc-MAPP-3L and imaging at 5 dpi, we mapped sucrose exudation across four eudicot (*A. thaliana*, *Brassica rapa*, *Brassica napus*, *Solanum pennellii*), and three monocot (*Sorghum bicolor*, *Setaria italica*, and *Setaria viridis*) species. Despite similar age and primary root length, monocot members exhibited lower biosensor signals along their entire primary root compared to eudicot members, with *B. napus* having a notably high biosensor signal at the root tip (Figure 2f). Bacterial colonization quantified by mScarlet fluorescence showed qualitatively similar spatial patterns across most roots, with average abundances that did not correlate with plant taxa, indicating that biosensor signals reflect exudation differences and that Suc-MAPP generally functions in species beyond *A. thaliana*.

To further validate phylogenetic differences and orthogonally quantify spatial exudation patterns, we adapted a previously reported in-media biosensor protocol that separates exudate detection from biosensor colonization^50^. We imaged *A. thaliana*, *S. viridis*, and *S. italica* seedlings transferred to agar diffusely embedded with *P. putida* Suc-MAPP-3L and observed exudate diffusion away from root tissues (Figure 2g). This assay corroborated root tip-inoculation results where *A. thaliana* signal intensities were highest around mature tissue and low at the root tip, while *S. viridis* and *S. italica* signals were overall low (Figure S11). Collectively, these data establish bacterial sucrose biosensors as robust tools for spatially profiling rhizosphere exudation, revealing previously uncharacterized exudation patterns across plant taxa.

### Biosensors Link Source–Sink Strength to Sucrose Exudation

We next used our *P. putida* biosensors to interrogate regulators of sucrose exudation in *A. thaliana* roots. Given sucrose’s role as the primary photosynthetic product governing source–sink carbon allocation, where shoots export sucrose to non-photosynthetic root tissues, we hypothesized that perturbing this process would alter biosensor-detected exudation. Source strength (i.e., sucrose production rate) was modulated by growing *A. thaliana* Col-0 under varying light regimes (ca. 20 to 100 μmol m^−2^ s^−1^). Decreasing photon flux led to reduced sucrose exudation, as measured by Suc-MAPP-3L in mature root regions (Figure 3a–b). In contrast, Suc-MAPP-3H signal was low and relatively undifferentiated between light conditions (Figure S12), consistent with its higher sucrose detection threshold (Figure 1f). Similar trends arose with chemical dampening of source strength via sub-lethal supplementation of the photosynthesis inhibitor and herbicide, atrazine^51^; atrazine-treated plants exhibited lower Suc-MAPP-3L-measured exudation across roots relative to atrazine-free controls (Figure 3c–d). Together, this suggests that sucrose exudation is gated by the rate of photosynthetic sucrose production and delivery to roots.

**Figure 3.**
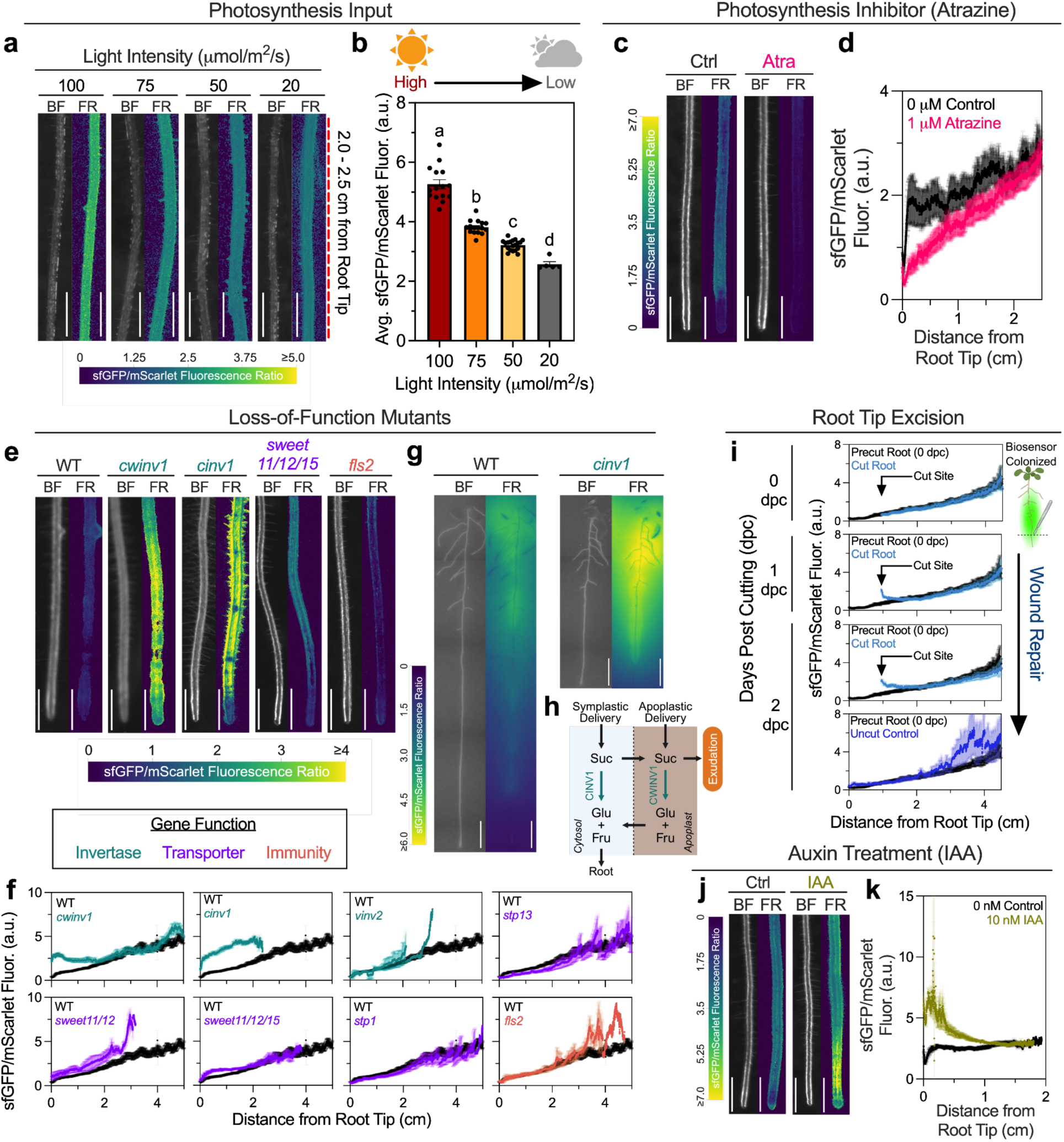
Sucrose Exudation is Modulated by Source–Sink Determinants. **a,** Representative images displaying sfGFP/mScarlet fluorescence along a midroot section (2.0-2.5 cm from root tip) of *A. thaliana* Col-0 seedlings grown at decreasing light intensity (100–20 µmol m^−2^ s^−1^). Roots were imaged 5 dpi with Suc-MAPP-3L biosensor. Scale bar, 1000 µm. **b,** Quantification of Suc-MAPP-3L biosensor signal on Col-0 grown under varying light intensity, averaged along whole roots (root tip to inoculation point). Significance groups were determined by one-way ANOVA with post-hoc Tukey’s, P < 0.05; letters correspond to statistically distinct groups. n = 15, 16, 19, 5 (from left to right) seedling replicates across 2 independent experiments. **c,d,** Sucrose exudation response to exogenous atrazine. Representative images displaying sfGFP/mScarlet fluorescence along a root tip section (0-0.5 cm) of *A. thaliana* seedlings Col-0 on solid medium supplemented with 1 µM atrazine or a DMSO control. Roots were imaged 5 dpi with Suc-MAPP-3L biosensor. Scale bar, 1000 µm. **d,** Quantification of sfGFP/mScarlet fluorescence (mean ± SEM) along whole roots. **e,f,** Quantification of sfGFP/mScarlet fluorescence along whole roots of *A. thaliana* sucrose metabolism knockout lines: invertases (teal), transporters (purple), and immunity (orange). Wild-type (WT) Col-0 (black) are shown in each graph for comparison. Roots were imaged 5 dpi with Suc-MAPP-3L *P. putida* biosensor. Scale bar, 1000 µm. **g,** Example images of 10 day old Col-0 (black) and *cinv1* (teal) *A. thaliana* seedlings imaged 24 hours post transfer to Suc-MAPP-3L-embedded solid medium. Scale bar, 0.5 cm. **h,** Model of cytoplasmic (CINV1) and cell wall (CWINV1) invertase-mediated effects on sucrose exudation. **i,** Measurement of sfGFP/mScarlet fluorescence (mean ± SEM) along the primary root of *A. thaliana* Col-0 seedlings 0-2 days post root tip (0-1 cm) excision relative to an uncut (blue) or the precut control (black). **j,k,** Sucrose exudation response to exogenous IAA. Representative images displaying sfGFP/mScarlet fluorescence along a root tip section (0-0.5 cm) of *A. thaliana* Col-0 seedlings on solid medium supplemented with 10 nM IAA or a DMSO control. Roots were imaged 2 dpi with Suc-MAPP-3L biosensor. Scale bar, 1000 µm. **k,** Quantification of sfGFP/mScarlet fluorescence (mean ± SEM) along whole roots.

To determine whether tissue sink strength (i.e., sucrose utilization rate) similarly constrains rhizosphere sucrose availability, we profiled a panel of *A. thaliana* T-DNA insertion mutants defective in key sink-determining pathways, including sucrose-hydrolyzing invertases^52–54^ and sugar transporters^55,56^. Applying Suc-MAPP-3L via the root tip-inoculation protocol, loss of cell wall invertase (CWINV1) or cytoplasmic invertase (CINV1) caused a pronounced increase in sucrose biosensor signal relative to the wild-type Col-0 control (WT) (Figure 3e–f). Rhizosphere sucrose levels from *cwinv1* appeared higher at young root tissues (0–1 cm from root tip), while *cinv1* had high exudate release across the entire root. Biosensor signal increases were not attributable to differences in bacterial abundance, as *cwinv1* exhibited colonization comparable to WT, while *cinv1* displayed regions of both elevated and reduced colonization relative to WT (Figure S13). The *cinv1* exudation phenotype was orthogonally confirmed via the in-media Suc-MAPP-3L biosensor assay (Figure 3g, Figure S11). In contrast, loss of vacuolar invertase (VINV2), sucrose transporters (SWEET11, SWEET12, and SWEET15), and hexose transporters (STP1, STP13) produced minimal changes in biosensor activity across young root tissues. To test whether presence of the *P. putida* biosensor triggers immunity-related alterations to host sink strength and spatial exudation responses, we also analyzed a mutant of the bacterial-recognition immune receptor FLS2^57^ and similarly observed no change in exudation at young root tissues (Figure 3e–f). As CWINV1 and CINV1 are both expressed at *A. thaliana* root tips^53,58^, biosensor activation in this region with knockout lines identifies apoplastic and cytosolic sucrose hydrolysis as major regulators of root sugar efflux. This indicates that metabolic consumption, rather than transporter capacity, governs exudation (Figure 3h).

Because plant tissue damage perturbs source–sink dynamics, we next asked whether injury impacts sucrose exudation. Root tip wounding increases local sink strength through elevated cell wall invertase expression and auxin signaling^59,60^, promoting apoplastic sucrose remobilization to support repairing tissue. Consistent with this framework, we observed a localized increase in Suc-MAPP-3L biosensor activity over two days in Col-0 roots excised ∼1 cm from the tip, relative to the same roots immediately prior to cutting (Precut Root) and non-excised controls over the same timescale (Uncut Control), indicating increased sucrose availability at the wound interface (Figure 3i). As root tip excision results in auxin pooling at the cut site^60^, we also tested whether auxin alone could cause a similar induction of exudation. Indeed, IAA supplementation to Col-0 led to a large sucrose biosensor signal accumulation within 0–0.5 cm of the root tip (Figure 3j–k). Given bidirectional cross-talk between sucrose and auxin, where sucrose delivery stimulates auxin biosynthesis^59^ and auxin reinforces sink-driven sucrose utilization^61^, our wounding and IAA supplementation findings suggest that sink pathways enhance sucrose exudation at wounded tissues.

### Synthetic Root Exudation Circuit Profiling by Biosensors

Genetically reprogramming root sucrose release could enable controlled dissection and manipulation of exudate effects on plant physiology and plant–microbe interactions. Thus, we applied a plant synthetic biology–bacterial biosensor approach to enhance sucrose levels at a low-exudation region in *A. thaliana*: the root tip (Figure 2). We hypothesized that cytosol-to-apoplast transport limits exudation, and that tunable, ectopic expression of SWEET11^62^ could quantitatively control sucrose release and activate *P. putida* biosensors. To test this, we engineered sets of SWEET11 Buffer Gate circuits^63^, where each set shares the same tissue-specific input promoter but the magnitude of SWEET11 expression output is controlled via a synthetic transcription factor (AmtR-ERF2-NLS) (Figure 4a). SWEET11 was translationally fused to the fluorescent reporter mTurquoise2, enabling quantitative mapping of local expression patterns while remaining optically compatible with sfGFP- and mScarlet-expressing *P. putida* biosensors (Figure S14).

**Figure 4.**
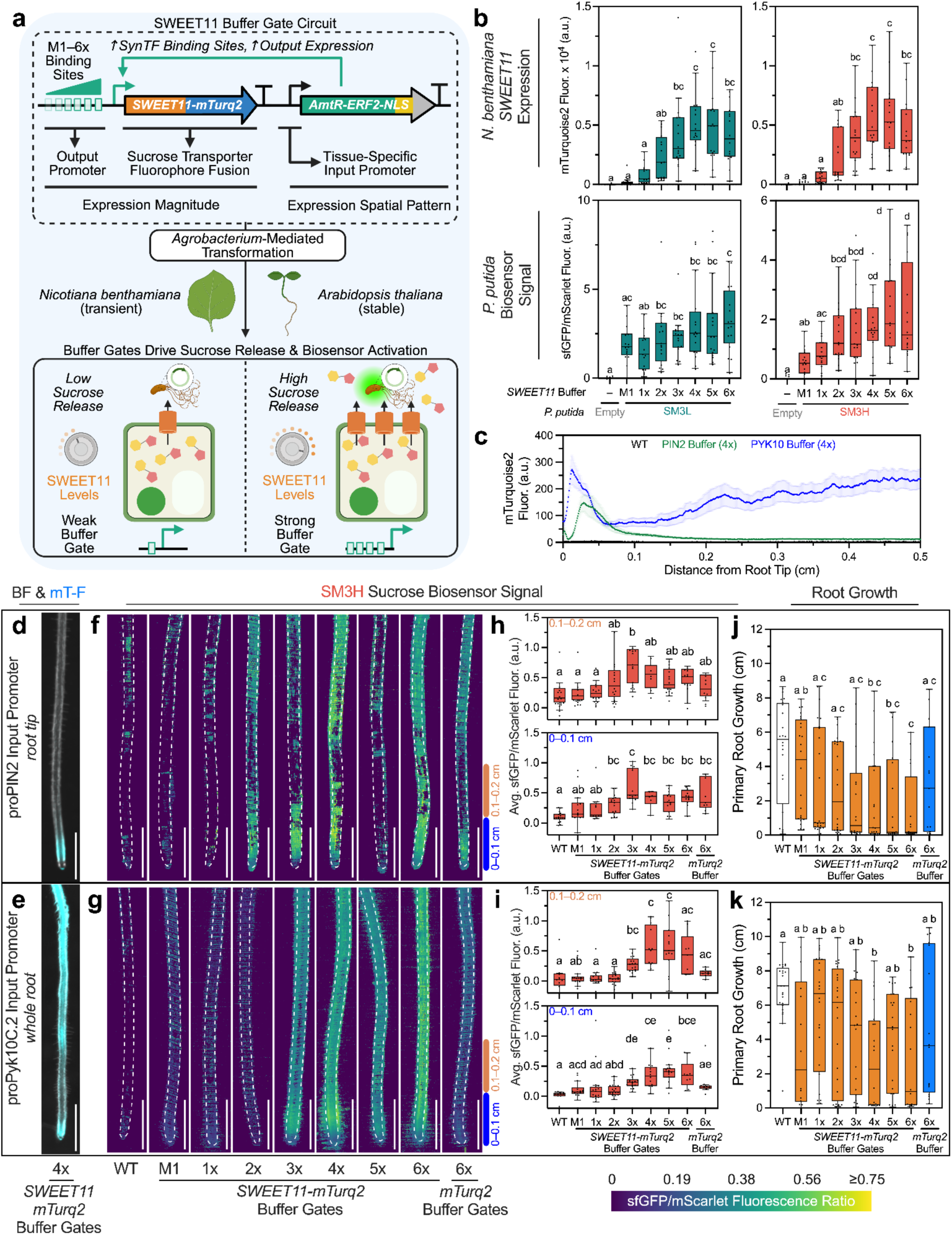
Sucrose Biosensors Phenotype Engineered Sucrose Exudation. **a,** SWEET11 Buffer Gate circuit design summary describing output promoter-driven expression magnitude and input promoter-driven spatial patterning along the root. Circuits were used to transiently (*N. benthamiana*) and stably (*A. thaliana*) engineer plant tissue to synthetically tune cellular SWEET11 expression and sucrose exudation. **b,** Quantification of mTurquoise2 fluorescence (mean ± SEM) of SWEET11-mTurq2 Buffer Gate *N. benthamiana* leaf discs then sfGFP/mScarlet fluorescence (mean ± SEM) post infiltration with either Suc-MAPP-3L (teal) and Suc-MAPP-3H (orange) biosensors. n = 11 (empty), n = 16 (M1–6x). Significance groups were determined by one-way ANOVA with post-hoc Tukey’s, P < 0.05 **c,** Quantification of mTurquoise2 fluorescence (mean ± SEM) along the primary roots of T3-generation *A. thaliana* 4x–*SWEET11*-*mTurq2* Buffer Gates driven by proPIN2 (green) or proPyk10C.2 (blue) input promoters. **d,e,** Representative bright field/mTurquoise overlay images of T3 *A. thaliana* 4x SWEET11 Buffer Gates driven by proPIN2 **(d)** and proPyk10C.2 **(e)** input promoters. Scale bar, 1000 µm. **f,g,** Representative images displaying sfGFP/mScarlet fluorescence along the primary roots of T1 *A. thaliana* Buffer Gates driven by proPIN2 **(f)** and proPyk10C.2 **(g)** input promoters imaged 5 dpi with Suc-MAPP-3H. Root traces from brightfield images are overlaid as dashed lines. Scale bar, 1000 µm. **h,i,** Quantification of average sfGFP/mScarlet across 1000 µm primary root sections, 0-0.1cm (blue) and 0.1-0.2 µm (orange). Significance groups were determined by Kruskal–Wallis with post-hoc Dunn’s, P < 0.05. **(h)** proPIN2, n = 18, 14, 14, 16, 14, 6, 15, 12, 9 (from left to right) seedling replicates across 2 biological replicates. **(i)** proPyk10C.2, n = 12, 14, 17, 18, 17, 12, 15, 9, 10 (from left to right) seedling replicates across 2 independent experiments. **j,k,** Primary root growth of T1 SWEET11 Buffer Gate seedlings post transfer to sucrose drop-out medium. Significance groups were determined by Kruskal–Wallis with post-hoc Dunn’s, P < 0.05. **j,** proPIN2, n = 24, 24, 24, 24, 18, 22, 20, 16, 15 (from left to right) seedling replicates across 2 independent experiments. **k,** proPyk10C.2, n = 24, 14, 24, 24, 19, 20, 22, 18, 16 (from left to right) seedling replicates across 2 independent experiments.

We prototyped constitutively-driven (proUBQ10 input) SWEET11 Buffer Gates via transient expression in *N. benthamiana*. Buffer Gate T-DNA was delivered to leaf tissue via *Agrobacterium tumefaciens* GV3101 infiltration, and subsequently co-infiltrated with either *P. putida* Suc-MAPP-3L or Suc-MAPP-3H biosensors. Subsequent fluorescence quantification confirmed that SWEET11 Buffer Gates dial transporter expression levels and differentially activate bacterial biosensors; Suc-MAPP-3H responded dose-dependently to SWEET11 expression strength, whereas Suc-MAPP-3L was uniformly activated across all Buffer Gates (Figure 4b). This suggests that Buffer Gates effectively tune apoplastic sucrose above the low micromolar sensing threshold of Suc-MAPP-3L, while staying within the sensing window of Suc-MAPP-3H (Figure 1f). These transient expression results highlight the utility of using biosensor bacteria for rapid phenotyping of engineered exudation circuitry.

We next tested root-expressing SWEET11 Buffer Gates in stable, transgenic *A. thaliana* lines at the T1 and T3 generations. The Buffer Gate tissue-specific input was swapped to proPIN2^62,64^ (root tip) or proPyk10C.2^65^ (whole root) promoters, with T3 mTurquoise fluorescence confirming the expected spatial patterns (Figure 4c–e; Figure S14). T1 lines were root-tip inoculated with the Suc-MAPP-3H biosensor, exhibiting graded biosensor activation at the root tip relative to wild-type Col-0 (WT) (Figure 4f–g). These results aligned with *N. benthamiana*–Buffer Gate experiments and the known Suc-MAPP-3H sensing threshold (Figure 1f); weaker SWEET11 expression (M1-2x) produces no measurable difference from WT, whereas stronger expression (3x-6x) is required to activate Suc-MAPP-3H (Figure 4h–i). These trends held across multiple independent transgene insertions (6–18 T1 seedlings tested per construct) and persisted at the homozygous T3 generation (Figure S15).

Interestingly, primary root growth inversely correlated with SWEET11 Buffer Gate strength across both root-specific promoter architectures (Figure 4j–k), with a greater inhibition by the proPIN2 input. This was largely driven by a subset of seedlings exhibiting root growth arrest post-transfer to sucrose-free media, which increased in frequency with Buffer Gate strength (Figure S16). Control lines expressing mTurquoise2-only Buffer Gates exhibited reduced, though not significantly different, growth relative to WT (Figure 4j–k). Together, these results suggest that enforced sucrose exudation and the transgene circuitry itself constrains root growth, likely by drawing on biomass-fueling carbon budgets and burdening transcription–translation resources^66^. Together, our *N. benthamiana* and *A. thaliana* results highlight that Buffer Gates and exudate biosensors enable quantitative trait engineering for root sucrose release, revealing physiological design rules and tradeoffs for synthetic exudation.

### Synthetic Community Abundances Track with Biosensor-Measured Sucrose

As proof-of-concept for directing rhizosphere ecology, we tested whether *A. thaliana* Buffer Gates could shift assembly of a 20-member synthetic bacterial community (SynCom)^67,68^. A pooled SynCom was applied to T3 proPIN2-SWEET11 Buffer Gate or Col-0 roots, and relative abundances at the root tip were measured by 16S rRNA profiling and regressed against parallel biosensor Suc-MAPP-3H measurements as a proxy for rhizosphere sucrose levels (Figure 5a–b). While most taxa were ‘non-responsive’, a subset of ‘responsive’ strains showed statistically significant correlations with rhizosphere sucrose levels, either positive (*Arthrobacter*, *Leifsonia*, *Ralstonia*, *Rhizobium*) or negative (*Agrobacterium* and *Bacillus*) (Figure 5c). Plant growth-enriched bacteria have been characterized by slower growth rates, which are associated with greater substrate utilization efficiencies in the rhizosphere^11^. Aligning with this, some of our responsive strains exhibit slower growth on sucrose opposed to non-responsive strains; while this trend was not statistically significant (sucrose P = 0.35; glucose P = 0.72), the effect was weaker on glucose (Figure S17–S18). These patterns suggest that (1) negative responders may be outcompeted, while (2) some positive responders may display stronger sucrose utilization efficiency, or (3) benefit from cross-feeding with non-responsive sucrose-catabolizers. Consistent with this, many non-responsive strains exhibited high sucrose growth rates and bioinformatically predicted sucrose metabolisms (Supplementary Table 1). Together, these results demonstrate that sucrose biosensors can quantify changes in environmental metabolites, including synthetically modulated exudation, and their effect on rhizosphere community shifts.

**Figure 5.**
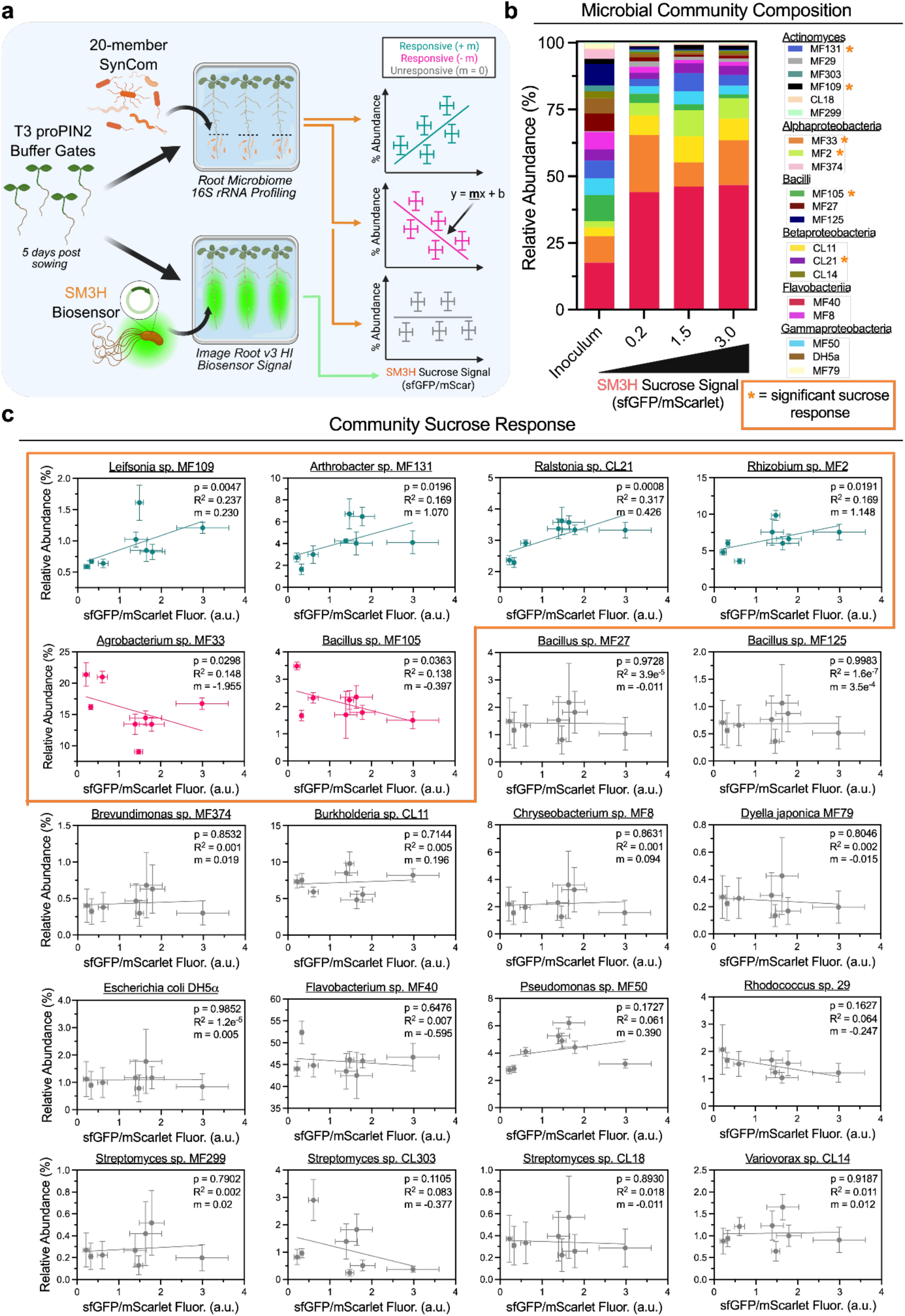
Programmed Sucrose Exudation Shifts Microbiome Assembly. **a,** Experimental outline highlighting the microbiome sucrose response assay. Either a 20-member SynCom or the Suc-MAPP-3H biosensor was inoculated on the roots of T3 proPIN2::SWEET11 Buffer Gates and 16S rRNA abundance was profiled at the root tip (0-1 cm). Significance (P < 0.05) between average strain abundance and average root tip v3-HI sfGFP/mScar ± SEM was determined via a simple linear regression. Slope sign (+/− m) identified correlation direction: sucrose responsive-positive (teal), responsive-negative (magenta), unresponsive (gray). **b,** Whole SynCom average strain abundance across T3 Buffer Gate-grown communities in order of increasing Suc-MAPP-3H (SM3H)-measured sucrose exudation. Root inoculum is provided as a reference. Strains are categorized taxonomically by class, and significant sucrose response (P < 0.05) is denoted by an orange asterisk. **c,** Changes in SynCom strain relative abundance (mean ± SEM, n = 4 replicates of pooled root tips) with increasing root tip sucrose exudation (mean sfGFP/mScar ± SEM, n = 12 seedling replicates) across the Buffer Gates.

## Discussion

Spatial heterogeneity of root exuded carbon underpins diverse rhizosphere processes^69,70^, yet inability to measure these gradients limits the study and manipulation of plant–microbe–environment interactions. By developing the Suc-MAPP suite of whole-cell sucrose biosensor circuits, we enabled root-associated *P. putida* to map previously unobserved exudation patterns across colonized plant tissues, dissect their environmental, physiological, and genetic drivers, and engineer localized release through synthetic exudation circuitry. Application of these biosensors across diverse plant rhizospheres, along with preliminary results in the *N. benthamiana* phyllosphere, suggests that whole-cell biosensing can be extended to quantify spatial gradients of additional metabolites identified through total exudate profiling^11,12^.

Although root tips are considered exudation hotspots^71,72^, our biosensor analysis across diverse plant species revealed that sucrose release at the primary root tip was generally low (Figure 2–3). However, exudation could be modulated by altering source–sink dynamics through physical (growth light intensity, tissue damage), chemical (photosynthesis inhibitors, auxin), or genetic perturbations (invertase knockouts) (Figure 3). The primary root^73^ and shoot meristems^74^ are characterized by high sink strength (sucrose demand) to support cell division and maintain apical dominance. Since intracellular sucrose levels in the root meristem are elevated relative to other developmental zones^14^, lower tip exudation likely reflects active conservation of this growth resource. Consistent with this interpretation, Buffer Gate circuits driving sucrose release at the root tip impaired root growth, particularly in the tip-localized proPIN2 variants (Figure 4j). Together, these results situate exudation within source–sink carbon budgets and set the stage for coupling biosensors with single-cell genomics to decode the genetic logic governing intra-versus extracellular carbon flux partitioning^75^.

Sucrose biosensor profiling of SWEET11 Buffer Gates demonstrates that engineered rhizobacteria can act as phenotypic outputs for plant synthetic biology and exudate engineering (Figure 4). While biosensors are extensively used to optimize metabolic pathways in unicellular systems^76,77^, their utility in multicellular plants is constrained because intact tissues cannot be analyzed using high-throughput technologies like FACS or microfluidics^78^. Our biosensor bacteria–transgenic plant system provides a hybrid approach in which root-colonizing biosensors enable non-destructive, fluorescence-based readouts of engineered traits in living plants, offering a scalable, high-throughput alternative to traditional metabolomics. Although demonstrated here for sucrose, this platform could be extended to monitor and engineer other plant-exuded bioproducts that influence microbial recruitment, nutrient mobilization, or plant immunity^8^.

Building on efforts to achieve high spatial resolution of microbiome biogeography^79,80^, our biosensor framework quantifies the root zone-scale influence of plant-released sucrose on rhizosphere community assembly (Figure 5). Our biosensor measurements were performed separate from synthetic community experiments, which may limit their accuracy in capturing sucrose levels in a community context. Future work could embed biosensor strains directly within communities to enable more precise *in situ* quantification of metabolite availability. Nonetheless, these measurements provide a previously inaccessible environmental variable that could help identify functional taxa responding to changes in sucrose exudation^81^, including those associated with developmental transitions^11,30^. Biosensors and Buffer Gate circuits could also be adapted to detect and modulate other spatially stratified rhizosphere metabolites, such as glutamine^6^, to probe their influence on microbial ecological functions like carbon source preference^82,83^ and chemotaxis^84^. By harnessing synthetic biology to investigate transkingdom metabolite exchange, Suc-MAPP biosensors offer a versatile platform to study and engineer plant–microbiome interactions for enhanced crop performance and sustainability.

## Methods

### DNA assembly and bacterial transformation

All bacterial strains, plasmids, and DNA sequences are listed in Supplementary Table S2. Plasmids were assembled via modular cloning Golden Gate assembly using BsmBI or BsaI (New England Biolabs) with DNA fragments generated by PCR (Phusion High-Fidelity, NEB) or commercially synthesized (Twist Bioscience). Reactions (10 μL) contained 25 fmol backbone and 50 fmol of each insert and were cycled 30–90 times depending on the number of parts (5 min at 37 °C for BsaI or 42 °C for BsmBI, followed by 5 min at 16 °C), then incubated at 60 °C for 10 min before storage at 10 °C. Golden Gate reactions were used to transform freshly prepared electrocompetent *Escherichia coli* and *Pseudomonas putida*. Purified plasmids were used to transform electrocompetent *Agrobacterium tumefaciens*.

Electrocompetent *P. putida* was prepared from glycerol stocks by overnight growth in LB at 28 °C, washing three times with 10% glycerol, and concentrating to ∼300 μL. Golden Gate reactions (0.5 μL) were mixed with 30 μL cells and electroporated at 1250 V. Cells were recovered in 250 μL LB for 2 h at 28 °C, plated on LB+25 μg mL⁻¹ kanamycin, and incubated overnight. Single colonies were expanded in LB+kanamycin, glycerol stocks prepared, and plasmids harvested for sequencing (Oxford Nanopore, Plasmidsaurus). Electrocompetent *A. tumefaciens* GV3101+pSOUP was prepared similarly, with initial growth and recovery in LB containing gentamicin (50 μg mL⁻¹) and tetracycline (10 μg mL⁻¹), and post-electroporation selection on LB agar containing gentamicin, tetracycline, and kanamycin (50 μg mL⁻¹). Colonies were expanded and stored as glycerol stocks.

### Sucrose biosensor construction and *in vitro* fluorescence assays

To identify sucrose-responsive biosensor circuits designs, we screened a library of plasmid variants in *P. putida* in which promoter (Figure S2) and ribosome binding site strengths^85–87^ were varied to tune expression of a sucrose sensor operon (Figure S3). The operon consisted of a LacI-family sucrose-responsive repressor (SuxR) from *Xanthomonas campestris*^44^ and outer (CscY) and inner membrane (CscB) sucrose transporters from *Pseudomonas protegens* Pf-5^43^. A synthetic SuxR-regulated promoter (P_sux2_) was constructed by adapting the P_LlacO-1_ promoter^88^, with operator sequences identified by RegPrecise^89^ and Snowprint^90^ placed upstream of the –35 element and between the –35/–10 spacer region. Constructs were assembled on kanamycin-resistant, high-copy RK2-origin plasmids^39^, containing a constitutive^91^ reporter cassette (P_ECrrnB(disc)_::*mScarlet-I*) for sfGFP normalization and a *par* locus for *in planta* plasmid stability^92^.

The pooled library was generated by Golden Gate cloning, electroporated into *P. putida*, and single colonies were cultured overnight in LB+25 μg mL^−1^ kanamycin at 28 °C and 250 rpm. Library members, along with Empty Plasmid and P_sux2_::*sfgfp* control strains, were screened by subculturing 100-fold into 1x M9 medium+20 mM glucose+25 μg mL^−1^ kanamycin ± 1 mM IPTG for 24 h, followed by endpoint fluorescence measurements using a Tecan Spark platereader (sfGFP: excitation 488 nm, emission 508 nm, 5 nm bandwidth; mScarlet-I: excitation 569 nm, emission 593 nm, 5 nm bandwidth). A top performing variant (Suc-MAPP-1) was isolated, sequence-verified, and reassembled into a fresh *P. putida* background to eliminate potential secondary mutations arising from library screening and confirm sucrose-inducible activation (Figure 1c).

To optimize dynamic range and sensing window, we generated a series of biosensor variants derived from Suc-MAPP-1. P_sux2_ was modified by strengthening its –35/–10 elements with those from P_J23101_ and reducing SuxR operator sites to a single site after the –10 element, generating P_sux1_, and circuit Suc-MAPP-2 (Figure 1d). Because CscB/CscY transporters exhibit millimolar affinity, mismatched to the micromolar sensing range of SuxR, they were replaced with *Xanthomonas* SuxCA transporters to generate Suc-MAPP-3L (Figure 1e). Finally, substituting SuxR with LacI-family repressor CscR from *Pseudomonas protegens* Pf-5 and pairing it with a cognate output promoter (P_csc1_) produced Suc-MAPP-3H, shifting the sensing threshold to higher sucrose concentrations (Figure 1f).

For all variants, sucrose–fluorescence response functions were generated by first inoculating strains from −80 °C glycerol stocks into LB+25 μg mL^−1^ kanamycin and incubating at 28 °C and 250 rpm for 24 h. Cultures were then subcultured 100-fold into 1x M9 medium+20 mM glucose+25 μg mL^−1^ kanamycin and grown for an additional 24 h. Following pregrowth, cultures were OD-normalizing to an initial OD600 of 0.02 in 1x M9 medium+20 mM glucose+25 μg mL^−1^ kanamycin supplemented with defined sucrose concentrations and incubated for 24 h at 28 °C and 250 rpm. Endpoint fluorescence measurements were obtained by diluting cultures 50- to 200-fold in black-walled, clear-bottom 96-well plates (Greiner Bio-one #655906) and analyzed using a Tecan Spark platereader (sfGFP: excitation 488 nm, emission 508 nm, 5 nm bandwidth; mScarlet-I: excitation 569 nm, emission 593 nm, 5 nm bandwidth). Response function data was fit to the following four-parameter activating Hill function (Graphpad Prism Version 11.0), where *y* is the measured sfGFP/mScarlet fluorescence ratio, *I* is the supplemented inducer concentration, *min* is the fitted uninduced sfGFP/mScarlet fluorescence ratio, *max* is the fitted induced sfGFP/mScarlet fluorescence ratio, *K*_1/2_ is the fitted half-maximal activation concentration, and *n* is the fitted cooperativity parameter:

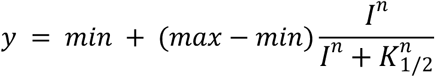

Dynamic range (DR) was defined as *max*/*min*. For the biosensor specificity experiment, cultures were prepared as described above except that the final 1x M9 medium+20 mM glucose+25 μg mL^−1^ kanamycin either contained either no carbon, 1 mM sucrose, or 1 mM of non-specific sugars (isomaltulose, trehalose, maltose, fructose, xylose). Additionally, fluorescence was measured after 6 h to minimize impacts to biosensing from sugar catabolism.

### *Nicotiana benthamiana*–sucrose biosensor infiltration assays

To assay sucrose biosensor induction in the *Nicotiana benthamiana* leaf apoplast, *P. putida* biosensors were inoculated from 25% glycerol stocks stored at −80 °C into 5 mL LB medium supplemented with 25 μg mL^−1^ kanamycin. The cultures were grown for 18 h at 28 °C and 250 rpm. 1 mL of culture was spun down (3,000 rcf, 10 min), washed three times with 10 mM MgSO4, and diluted to a final OD_600_ = 0.2. Biosensors were injected into previous *Agrobacterium* infiltration sites on the abaxial side of *N. benthamiana* leaves using a needleless syringe. Plants were returned to normal growth conditions consisting of 12 h light (25 °C, 75 μmol m^−2^ s^−1^)/12 h dark (22 °C) for 24 h. 8 mm discs at the infiltration sites were excised with a hole punch, leaf discs were placed in flat-bottom 96-well plates, and fluorescence was quantified using a plate reader in bottom-reading mode (Tecan SPARK).

*N. benthamiana* leaf infiltration was adapted from a previous protocol^63^. For *Agrobacterium tumefaciens* GV3101+pSOUP carrying binary vector plasmids or *Agrobacterium tumefaciens* GV3101+p19 was inoculated from 25% glycerol stocks at −80 °C into 2 mL LB+50 μg mL^−1^ kanamycin+50 μg mL^−1^ gentamicin+10 μg mL^−1^ tetracycline or LB+50 μg mL^−1^ kanamycin, respectively. The cultures were grown for 24 h at 28 °C and 250 rpm. Cultures were normalized to OD_600_= 0.02 using fresh LB medium with kanamycin then grown for another 24 h under the same conditions. Following growth, the cultures were spun down (3,000 rcf, 8 min) and resuspended in 2 mL of infiltration buffer (10 mM MgCl_2_, 10 mM MES pH 5.7, 150 μM acetosyringone) to incubate at room temperature for 3 h. Each final sample was a 1 mL mixed culture of target *Agrobacterium* strains normalized to OD_600_ = 0.3 and P19-silencing suppressor strains normalized to OD_600_= 0.2 using infiltration buffer. Samples were injected into the abaxial side of *N. benthamiana* leaves using a needleless syringe and returned to normal growth conditions for 48 h. After 48 h, *P. putida* strains were infiltrated into pre-agroinfiltrated spots as described above, returned to normal growth conditions for another 24 h, then fluorescence from 8 mm leaf discs were analyzed by a Tecan Spark platereader. All *N. benthamiana* agrofiniltrated T-DNA constructs expressing *SWEET11*-mTurquoise2 translational fusions contained an intron within the fluorescent reporter to prevent spurious fluorescence from bacterial expression^63^.

### Plant materials and growth conditions

*Arabidopsis thaliana* Col-0 ecotype was used throughout the study as a wild-type line and parent to Buffer Gate transgenic lines. Seed materials from *Brassica napus* (Darmor-bzh)*, Brassica rapa* (Yubileinaya Zelenogolovaya, PI 633171)*, Setaria italica* (GE.2013-27, PI 690671)*, Setaria viridis* (A10.1, WV001-9), and *Solanum pennellii* (LA1272, PI 365970) were obtained from the National Plant Germplasm System (NPGS), USDA-ARS, via the GRIN-Global database. *Sorghum biocolor* BTx623 seeds were obtained from Katrien Devos at the University of Georgia, Athens, USA. *A. thaliana* T-DNA knockout mutants *cinv1* (SALK_095807), *cwinv1* (CS25124), *vinv2* (SALK_100813), *stp1* (SALK_048848), and *stp13* (SALK_021204) were obtained from the Arabidopsis Biological Resource Center (ABRC) at Ohio State University, Columbus, OH. The *A. thaliana sweet11/12* and *sweet11/12/15* mutants were shared by Wolf B. Frommer, and *fls2* seeds were shared by Libo Shan. Eudicots were bulked for seeds or floral dip in a Conviron MTP144 at 150–180 μmol m^−2^ s^−1^ under 16 h light (22 °C) / 8 h dark (20 °C) cycles, in PRO-MIX HP Mycorrhizae soil (Pro-Mix 20381RG). Monocots were bulked for seeds in greenhouse conditions.

For biosensor phenotyping experiments, *A. thaliana* seedlings were grown on MS medium petri dishes at 100 μmol m^−2^ s^−1^ in a Conviron Plant Growth Chamber (Gen1000) at the University of Georgia with LED lighting at the University of Georgia, except Figure 2d–e and Figure 4d–k that were performed at 100 μmol m^−2^ s^−1^ in a Percival-Scientific Growth (CU36L4) at Stanford University with fluorescent lighting. Figure 3a–b was also performed in the Percival-Scientific Growth unit, but with chamber settings dialed to four distinct light intensities (20, 50, 75, 100 μmol m^−2^ s^−1^).

### Root-tip inoculation protocol

*Arabidopsis thaliana*, *Brassica napus*, *Brassica rapa*, and *Solanum pennelli*, *Sorghum bicolor*, were sterilized for 5 min with 70% EtOH, followed by 5 min with 20% bleach, then washed 3 times with sterile water. *Setaria italica* and *S. viridis* seeds were sterilized for 5 min only with 20% bleach + 0.04% Tween20. All seeds were cold stratified (4 °C) in sterile water 2 d prior to sowing.

Stratified seeds were sown on sucrose-supplemented plant medium consisting of ½ strength MS (Carolina,195703) + 1% sucrose + 0.7% gelzan in square petri dishes and grown vertically under a 16 h light (22 °C, 100 μmol m^−2^ s^−1^) / 8 h dark (20 °C) photoperiod for five days prior to their transfer onto sucrose drop-out MS medium. When required, we supplemented sucrose-dropout MS medium with chemical stressors: atrazine (1 µM), IAA (10 nM), and included DMSO-supplemented controls for all nonpolar chemical additives.

From 25% glycerol stocks at −80 °C, Suc-MAPP-3L and Suc-MAPP-3H *P. putida* biosensors were inoculated into 5 mL LB (Sigma-Aldrich, L3022)+25 μg mL⁻¹ kanamycin. The cultures were grown for 16 h at 28 °C and 250 rpm. The liquid culture was reinoculated into 5 mL 1M M9+glucose medium (5x M9 salts [Becton Dickinson, 248510] (200mL/L), 1 mM MgSO_4_, 0.1 mM CaCl_2_, 10 mM glucose)+25 µg µL^−1^ kanamycin to grow for another 16 h. The culture was spun down (3,000 rcf, 10 min), washed three times with 10 mM MgSO_4_, and diluted to a final OD_600_= 0.003. Seedling root tips were inoculated with 3 µL of diluted culture immediately following their transfer to MS-sucrose medium.

### IPTG-induced *lac* biosensor assay

*A. thaliana* seedlings were grown as described above for 5 days on sucrose-supplemented plant medium. Upon transfer to sucrose drop-out medium supplemented with either 0 mM or 1 mM IPTG inducer, seedling root tips were inoculated with 3 µL of OD_600_= 0.003 *lac* biosensor or control strains (grown under conditions identical to Suc-MAPP-3H/LOW sensors). To measure biosensor signal turn off, seedlings were transferred to fresh 0 mM IPTG medium after 5 d, then imaged 0, 1, 2, and 4 days post transfer.

### Root image analysis

Seedlings were imaged 2, 5, and 7 days post transfer to MS-sucrose medium using a Leica THUNDER Imager Model Organism with fluorescence filters either at the University of Georgia, or for Figure 2d–e, Figure 3a–b, and Figure 4d–k at Stanford University. For all experiments except T1-generation Buffer Gate experiments (Figure 4d–k), raw fluorescence pixel intensity (sfGFP, mScarlet, or mTurquoise channels) was quantified in Fiji (ImageJ 2.16.0/1.54p)^93^ using the Segmented Line tool and Plot Profile. A segmented line width of 20 was used for all analyses except the T1 Buffer Gate experiments due to minimal between-replicate variation and to match line width to root diameter (Figure S19). T1 Buffer Gates fluorescences were measured using 45pt segmented line width due to higher magnification. Average background fluorescence was determined from a plant-free region and subtracted from all raw intensity values. Along the primary root, sfGFP intensity was normalized to mScarlet at each position, producing a ratiometric sfGFP/mScarlet fluorescence signal that was plotted as a function of distance from the root tip. Representative fluorescence ratio images were generated in FIJI by dividing background-subtracted sfGFP by mScarlet channels. Global thresholds were applied such that only pixels with above-background mScarlet signal were retained, while background pixels were set to ratio of zero. The *mpl-viridis* lookup table was applied to the resulting ratio images.

### In-media biosensor assay protocol

The Suc-MAPP-3L agar preparation was adapted from a previous protocol^50^. The Suc-MAPP-3L biosensor was prepared as previously described. The final culture was spun down (3,000 rpm, 10 min) and cells were washed three times with 10 mM MgSO_4_. Cells were then resuspended to OD_600_= 0.45 in 1M M9+glucose medium.

The Suc-MAPP-3L bacterial biosensor plates consisted of the modified M9 glucose medium+1.6% Agar (BP9744500, Fisher Scientific; 16g/liter), then cooled to 45°C, and combined with 10% (v/v) of OD_600_ = 0.45 of Suc-MAPP-3L culture. After mixing, 50 mL of the biosensor agar medium was poured into 120-mm square Petri dishes and left to cool completely.

Wild-type (*A. thaliana* Col-0, *S. italica*, *S. viridis*) and *A. thaliana* mutant (*cinv1, cwinv1,* and T3 proPIN2::SWEET11 2x) seedlings were grown as previously described. After 4 days growth on MS-sucrose medium, they were transferred to Suc-MAPP-3L agar with sucrose standards consisting of sterile filter paper strips (5 mm x 40 mm) soaked in aqueous sucrose solutions (0, 100, or 1000 mM). Whole root fluorescence was imaged 24 h post transfer.

### Root excision assay

*A. thaliana* Col-0 seedlings were grown as previously described. Seedlings were inoculated with the *P. putida* Suc-MAPP-3L biosensor upon transfer to MS-sucrose medium, and precut root fluorescence was captured 5 days post inoculation. We excised 1 cm of root tip tissue from precut roots then imaged the fluorescence from the new root tips at 5, 6, and 7 dpi.

### Transgenic *A. thaliana* Buffer Gate construction

Transgenic *Arabidopsis thaliana* Col-0 plants were generated via *Agrobacterium tumefaciens*–mediated floral dip. Briefly, *A. tumefaciens* cultures were initiated from frozen stocks into 14-mL culture tubes containing 2 mL LB with appropriate antibiotics and grown for 24 h at 28 °C and 250 rpm. Following growth, 250 μL of each culture was plated on 15-cm LB agar plates with antibiotics and incubated for 48 h at 28 °C. Cells were then scraped from plates and resuspended in 200 mL transformation buffer (per liter: 50 g sucrose, 0.93 g MgCl₂, 250 μL Silwet L-77). Inflorescences were submerged in the bacterial suspension for 2 min, then incubated overnight in a dark, humid container. Plants were returned to standard growth conditions and grown to seed. Transgenic seeds were selected using the seed-specific RUBY marker (proAt2S3::RUBY), allowing identification of T1 plants via red pigment in dry seeds by naked eye, as described previously^94^.

### SynCom culture and plant inoculation

The 20-member SynCom strains were inoculated from glycerol stocks into 200 µL LB medium in a 96-well plate. Cultures were grown at 28°C while shaking at 270 rpm. After 48 h growth, cultures were OD_600_ normalized to each other, then pooled in equal volumes. The mixed culture was spun down (3,000 rpm, 2 min), washed three times with 10 mM MgSO_4_, then resuspended to a final OD_600_ = 0.2. T3 proPIN2 Buffer Gate::*SWEET11*–mTurq2 Buffer Gates (M1–6x) root tips were inoculated with 3 µL of pooled SynCom culture immediately following their transfer to solid MS-sucrose medium.

### Bacterial DNA extraction and 16S metagenomics

After 5 days of growth, seedling root tips (0-1cm) were harvested, pooling 6 roots for each plant genotype in quadruplicate. Roots were placed in a 2-mL PowerBead Pro Tube and stored at - 80°C until processing. Samples were homogenized using the Tissuelyser III (Qiagen, #9003240) and rhizosphere DNA was extracted using the PowerSoil Pro Kit (Qiagen, #47014), following the manufacturer’s instructions.

We amplified 16S rRNA using V3-V4 region primers with Illumina overhang adapters: forward (5’-TCGTCGGCAGCGTCAGATGTGTATAAGAGACAGCCTACGGGNGGCWGCAG-3’) and reverse (5’-GTCTCGTGGGCTCGGAGATGTGTATAAGAGACAGGACTACHVGGGTATCTAATCC-3’)^95^.

First round PCR reactions were carried out in 25 µL volumes containing 12.5 µL 2x KAPA HiFi HotStart ReadyMix (Roche), 0.75 µL of each 10 µM primer, and 2 µL template DNA. First round PCR conditions were as follows: 95°C for 3min; 30 cycles of 95°C for 30s, 62°C for 15s, 72°C for 30s; 72°C for 3 min; 4°C hold. The following 16S library preparation was adapted from a previous protocol^95^ with modifications. Then, 2 µL of the first round PCR product was diluted in 78 µL nuclease-free water (Fisher Bioreagents, BP2484100). We ligated i7/i5 indices and sequencing adapters to diluted PCR1 products using Nextera™-Compatible Multiplex Primers (Active Motif, 53155). Second round PCR reactions were carried out in 25 µL volumes containing 12.5 µL 2x KAPA HiFi HotStart ReadyMix, 2 µL of each 5 µM primer, and 2 µL template DNA. Second round PCR conditions were as follows: 98°C for 45 s; 9 cycles of 98°C for 15 s, 60°C for 30 s, 72°C for 30 s; 72°C for 60 s; 4°C hold. To purify, 5 µL of each round two PCR product was pooled and purified using the QIAquick PCR & Gel Cleanup Kit (Qiagen, #28506) following manufacturer’s instructions. The purified gel band was quantified using a Tecan Spark platereader and normalized to a final concentration of 10 nM. Sequencing was performed on an Illumina MiSeq instrument using a 500-cycle V2 kit (MS-102-2003).

### *In vitro* SynCom culture protocol

From glycerol stocks, SynCom strains were cultured in 5 mL LB at 28°C while shaking at 270 rpm. After 3 days, cultures were spun down (3,000 rpm for 10 min), washed three times with 10 mM MgSO_4_, and resuspended to OD_600_ = 1 in 10 mM MgSO_4_. In flat-bottom 0.2 mL 96-well plates, strains were reinoculated into 1M M9 medium supplemented with either glucose or sucrose in triplicate at a final OD_600_ = 0.02. Cultures were grown using a Tecan Spark platereader at 28°C while shaking at 270 rpm for 72 h, and OD_600_ was measured in 10 min intervals. From each strain’s OD_600_ growth curve, maximum growth rates (µ_max_) were calculated from the log phase, maximum yields (OD_600_max) from the stationary phase, and lag time was determined as the x-intercept of the line tangent to the µ_max_.

### 16S rRNA abundance analysis

All bioinformatic analyses were performed in R (version 4.5.2). Raw paired-end 16S rRNA gene amplicon files were processed using the DADA2 package^96^. Sequencing adapters and residual primer sequences were removed using cutadapt prior to downstream analyses. Quality filtered reads were processed to infer exact amplicon sequence variants. Error models were learned from the data, forward and reverse reads were merged, and chimeric sequences were identified and removed, resulting in an ASV count table and a table of representative sequences.

Taxonomic assignment was performed using a modified reference database based on SILVA v138 that included the exact 16S rRNA gene sequences of the SynCom members. Chloroplast-and mitochondria-derived ASVs were removed prior to downstream analyses. ASV count tables and taxonomy tables were synchronized to retain only taxa present in both datasets. Sample metadata were provided separately and included information on sample identifiers, plant genotype, treatment, replicate, and sample type. Metadata were curated to ensure consistency of sample identifiers and compatibility with downstream analyses.

ASV abundances were summarized at the species level for community composition analyses. Low abundance taxa were defined based on a relative abundance threshold of 0.1% across samples and were grouped into a single “low abundance” category for visualization purposes. (This threshold was used only for plotting and did not involve removal of taxa from the underlying ASV tables.)

All plots were generated in GraphPad Prism v10.6.0. A simple linear regression was performed between strain percent abundance and average Suc-MAPP-3H-determined sucrose at the root tip (0-1cm) to establish sucrose-responsive (P < 0.05) and -unresponsive (P > 0.05) strains.

## Acknowledgements

We thank members of the Dinneny, Dundas, and Wallace Labs for helpful input on the project, Benjamin K. Keitz for providing biosensor plasmid backbones, Rachel Porter and Kerwyn C. Huang for assistance with 16S rRNA profiling, Trent Northen for providing *Paraburkholderia graminis*, Jung-Gun Kim and Mary Beth Mudgett for providing *Xanthonomas* genomic DNA, Li Yang for providing SynCom strains, and the Georgia Genomics and Bioinformatics Core for performing 16S rRNA sequencing services. C.M.D was supported by a Tomkat Center Postdoctoral Fellowship in Sustainable Energy at Stanford University. G.A.B. was supported by a University of Georgia Research Institute Scholars’ Assistantship. J.R.D. is a Chan Zuckerberg Biohub—San Francisco Investigator and an investigator of the Howard Hughes Medical Institute.

## Author Information

### Contributions

C.M.D. and J.R.D. conceived the project. C.M.D., G.A.B., and J.R.D. designed research, analyzed data, and drafted the manuscript. C.M.D., G.A.B., T.C., M.P performed experiments.

J.A.U. and J.W assisted with microbiome analysis. I.V. assisted with Buffer Gate construct cloning and *Nicotiana* experiments. All authors gave input on the final manuscript.

## Supplementary Information

Supplementary Figures S1-S25.

