## Supplemental Information for "Rhizobacterial Biosensors Spatially Map Natural and Engineered Sucrose Exudation"

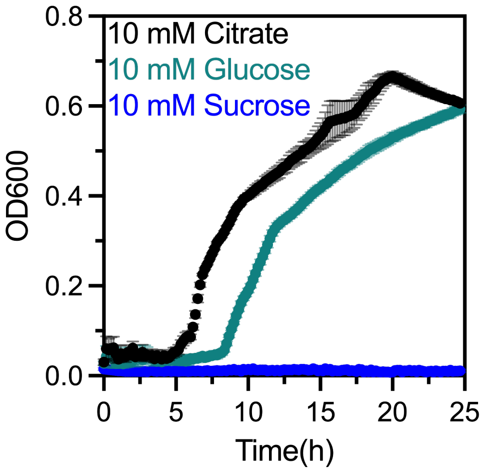


**Figure S1. Carbon source utilization in *Pseudomonas putida*.**
*P. putida* KT2440 (Empty Plasmid) growth, measured as optical density ($OD_{600}$), in liquid 1x M9+25 µg $mL^{-1}$ supplemented with either 10 mM citrate (black), 10 mM glucose (teal), or 10 mM sucrose (blue) (mean ± SEM, n=4 biological replicates).

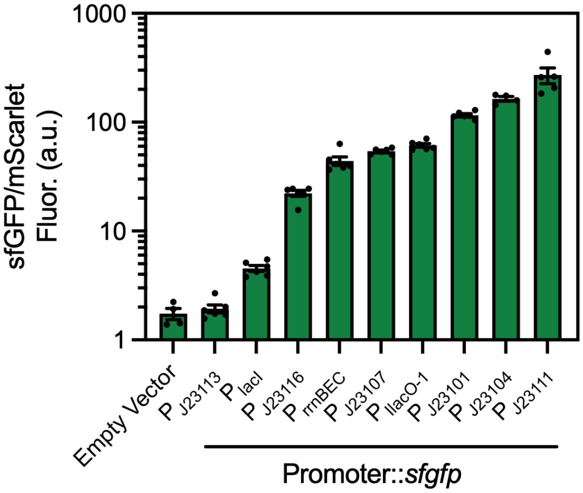


**Figure S2. Screening a promoter variant library in *Pseudomonas putida*.**

Ratiometric sfGFP/mScarlet fluorescence measured across a plasmid-expressed promoter::*sfgfp* variant library in *P. putida* KT2440. Strains were cultured in M9 minimal medium with 20 mM glucose for 24 hours prior to quantification (mean ± SEM, n=5 biological replicates).

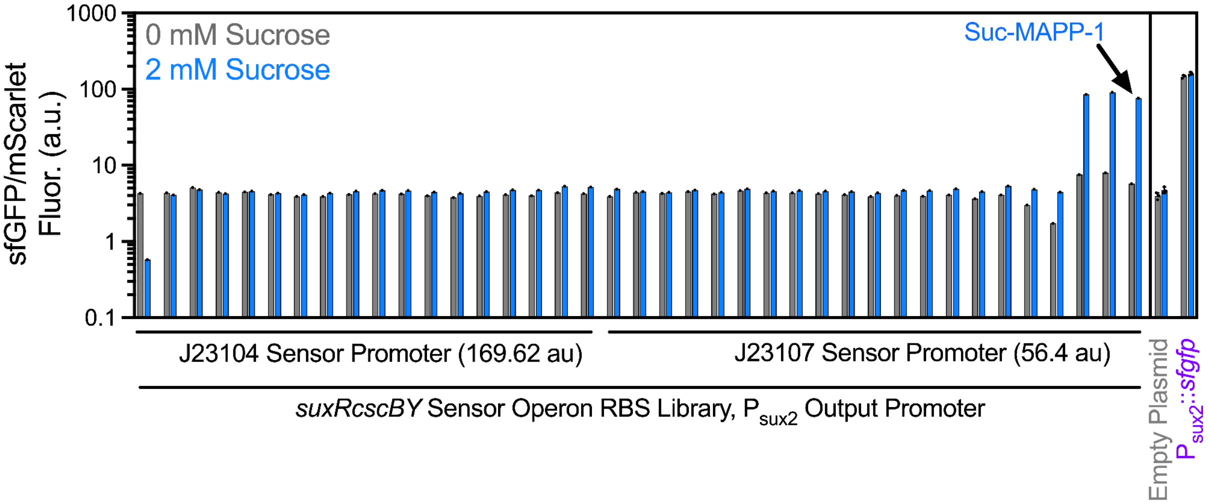


**Figure S3. Screening biosensor operon promoter/RBS library.**

Quantification of sfGFP/mScarlet fluorescence across a promoter and ribosome binding site (RBS) variant library for the biosensor operon genes (*suxR*, *cscB*, *cscY*). Promoter strengths (indicated in parentheses) were determined from sfGFP/mScarlet fluorescence ratios in Figure S2. RBS variants were designed using the DeNovoDNA RBSCalculator[2,3] with predicted strengths of: *suxR* (single RBS, 1134.84 a.u.), *cscB* (eight-member degenerate library, 3.61–1799.85 a.u.), *cscY* (eight-member degenerate library, 5.08–2058.81 a.u.). Libraries were assembled by one-pot combinatorial Golden Gate cloning of promoter–RBS variants driving biosensor genes, transformed into *P. putida* KT2440, and assayed as individually isolated library variants. Each bar represents a single variant grown under 24 h in 1x M9+20 mM glucose+25 µg mL$^{-1}$, under induced

(2 mM sucrose) or uninduced (0 mM sucrose) conditions. Negative (Empty Plasmid) and positive ($P_{sux2}$::*sfgfp*) control strains were included for comparison.

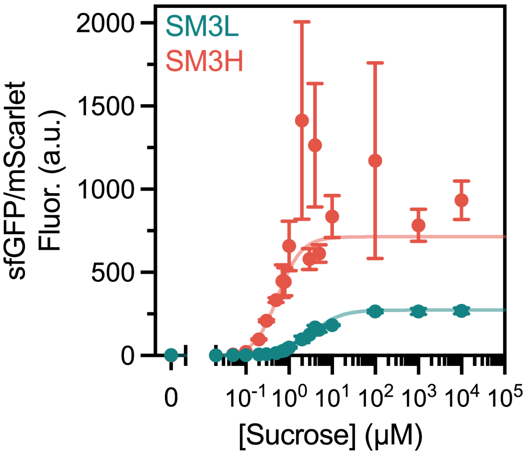


**Figure S4. Response function of Suc-MAPP biosensors deployed in *Paraburkholderia graminis*.**

Ratiometric sfGFP/mScarlet signal of Suc-MAPP-3L (SM3L) and Suc-MAPP-3H (SM3H) deployed in *P. graminis*. Cultures were grown in liquid M9–glucose–kanamycin media supplemented with varying sucrose concentrations.

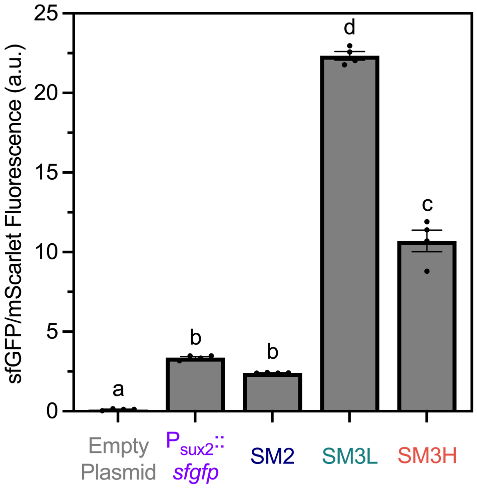


**Figure S5. *In vitro* induction of *Pseudomonas putida* biosensor strains in-parallel with *Nicotiana bethamiana* leaf infiltration.**

Ratiometric sfGFP/mScar signal (mean ± SEM) of Empty Plasmid, constitutive $P_{sux2}$::*sfgfp*, Suc-MAPP-2 (SM2), Suc-MAPP-3L (SM3L), and Suc-MAPP-3H (SM3H) circuits expressed in *P. putida.* All strains were grown in liquid M9 biosensor induction media supplemented with 1 mM sucrose.

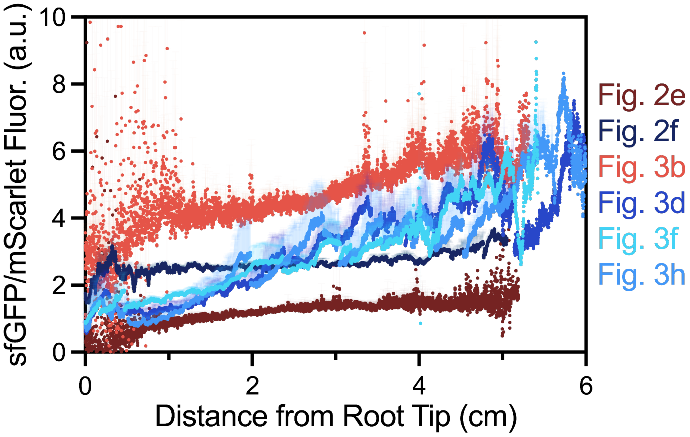


**Figure S6. *Arabidopsis thaliana* exudation meta-analysis across experiments.**
Ratiometric sfGFP/mScarlet fluorescence (mean ± SEM) measured along *A. thaliana* Col-0 roots at 5 days post inoculation with the *P. putida* Suc-MAPP-3L biosensor. Data were compiled from multiple experiments conducted using the same model fluorescence stereomicroscope (Leica M205 FCA) at either Stanford University (SU, orange) or the University of Georgia (UGA, blue). Plant growth at SU was performed with a Percival Scientific CU-36L4 at 16 h light/8 h dark cycle at constant temperature of 22 °C with light conditions corresponding to 100 µmol $m^2$ $s^{-1}$ of white fluorescent lights. Plant growth at UGA was performed with a Conviron Gen1000 at 16 h light (22 °C)/8 h dark (20 °C), with light conditions corresponding to 100 µmol $m^2$ $s^{-1}$ of LED lights. Datasets correspond to Fig. 2e (SU), Fig. 2f (UGA), Fig. 3b (SU), Fig. 3d (UGA), Fig. 3f (UGA), Fig. 3h (UGA).

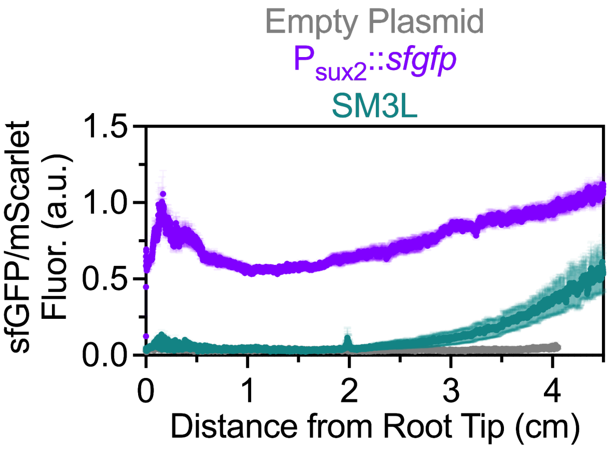


**Figure S7. Colonization of *Paraburkholderia graminis* on *Arabidopsis thaliana***
Ratiometric sfGFP/mScarlet signal (mean ± SEM) along *A. thaliana* Col-0 primary roots imaged at 5 days post inoculation with the sucrose biosensor (Suc-MAPP-3L/SM3L) and control (Empty Plasmid and $P_{sux2}$::*sfgfp*) *P. graminis* strains.

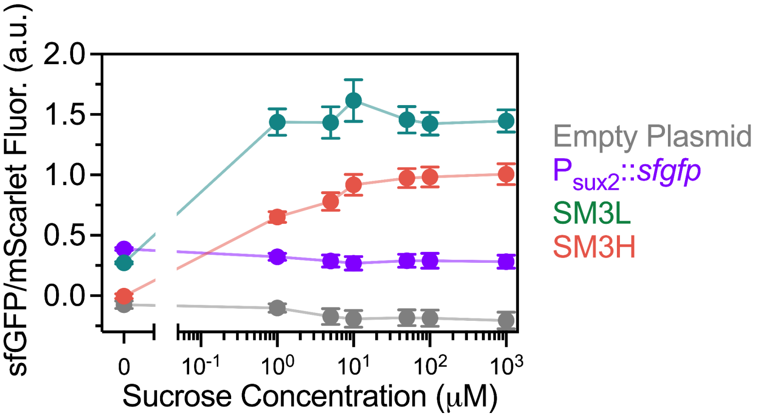


**Figure S8. *P. putida* biosensor activity on plant-free solid media.**
Ratiometric sfGFP/mScarlet fluorescence (mean ± SEM) of *P. putida* Suc-MAPP-3L (SM3L, teal) and Suc-MAPP-3L (SM3H, orange) biosensors imaged 5 days post inoculation on solid plant media (½ MS + 0.7% gelzan + 1 mM glucose) supplemented with sucrose (0-1000μM). $P_{Sux2}$::sfGFP (purple) and empty (teal) constructs were used as positive and negative controls, respectively.

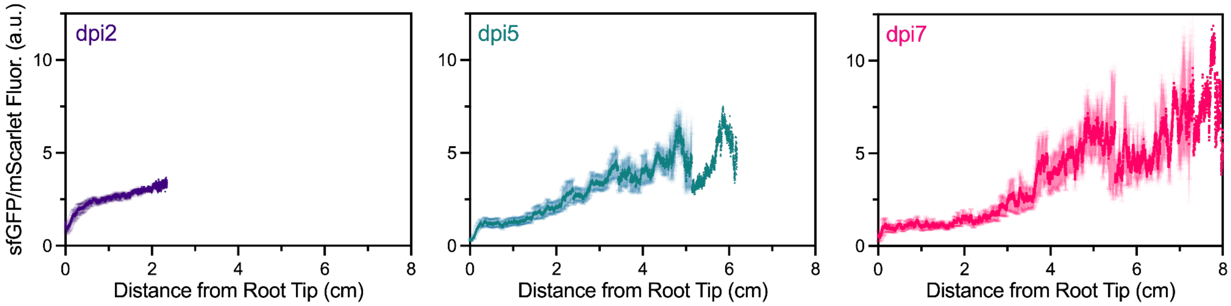


**Figure S9. Temporal analysis of sucrose biosensor activity on *Arabidopsis thaliana* roots.**
Ratiometric sfGFP/mScarlet fluorescence (mean ± SEM) along *A. thaliana* Col-0 roots imaged 2 (purple), 5 (teal), and 7 (magenta) days post inoculation with the *P. putida* Suc-MAPP-3L biosensor.

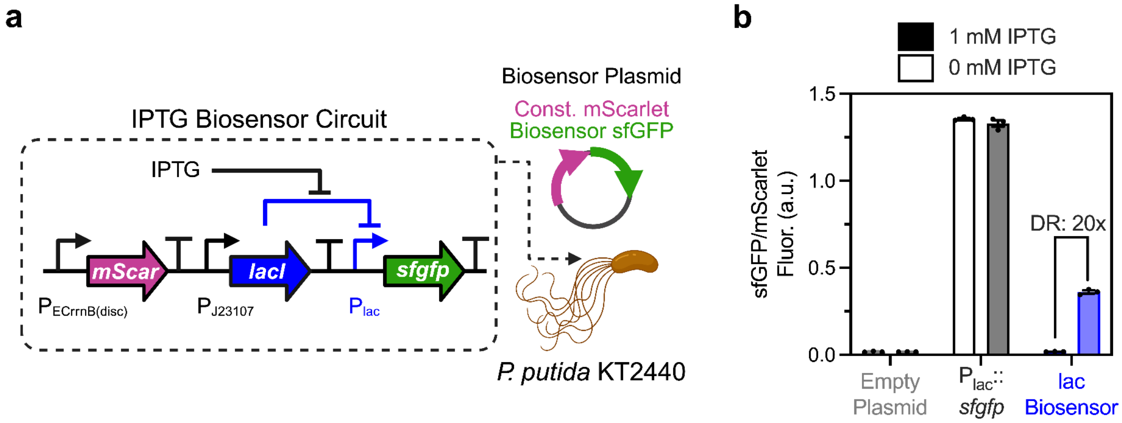


**Figure S10. Functional response of IPTG-responsive lac biosensor.**
**a,** Lac biosensor circuit design. **b,** Average ratiometric sfGFP/mScarlet signal ± SEM of the lac biosensor and control plasmids expressed in *P. putida* grown for 24 hours in liquid M9+20 mM glucose media supplemented with 0 or 1 mM IPTG. DR = Dynamic Range.

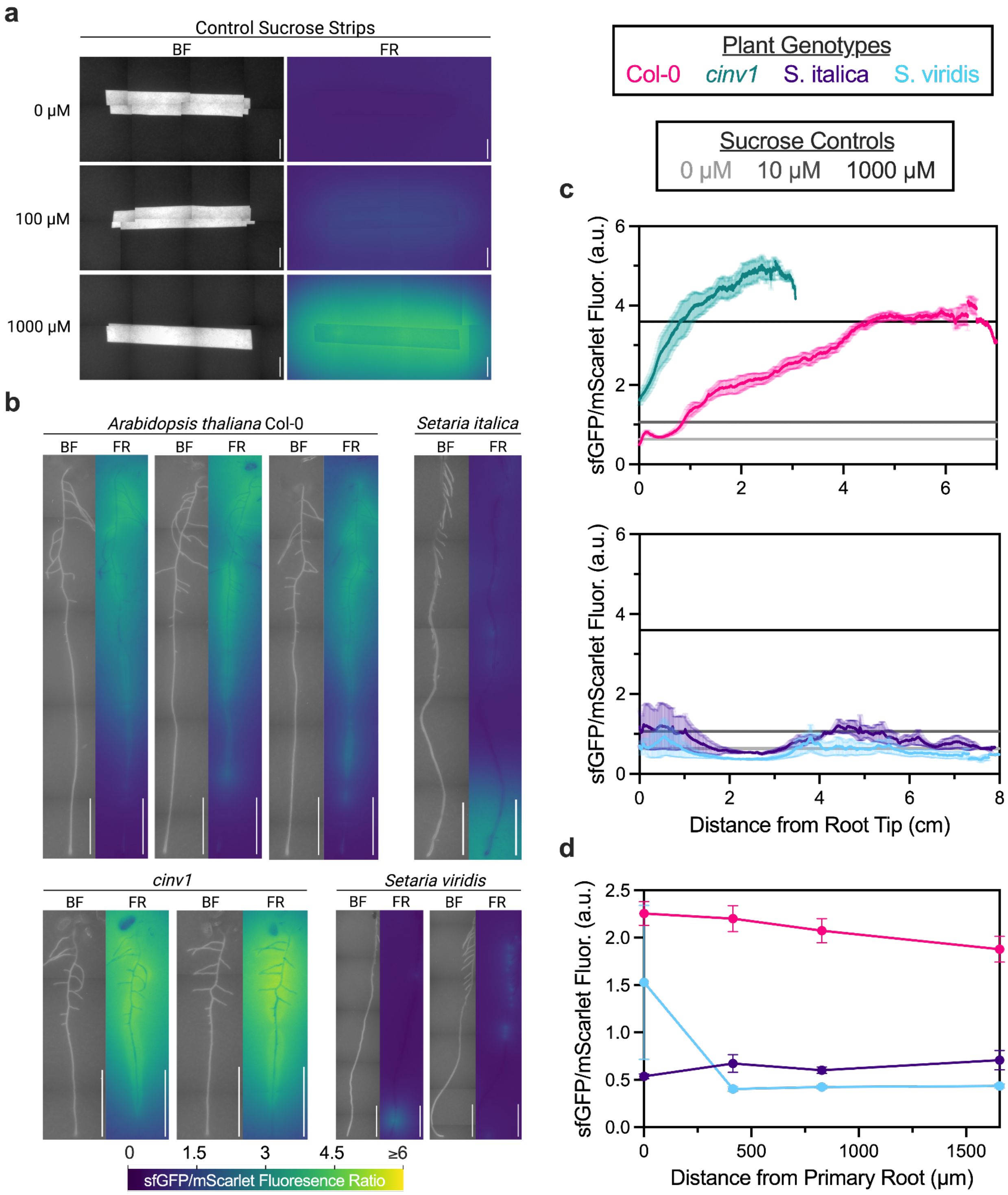


**Figure S11. Whole plant exudation profiles on biosensor-embedded agar.**
**a,b,** Ratiometric sfGFP/mScarlet fluorescence overlay images of sucrose filter paper strips **(a)** and whole plants **(b)** 24 hours post transfer to Suc-MAPP-3L-embedded agar. Scale = 5000 µm. **c,** Ratiometric sfGFP/mScarlet fluorescence (mean ± SEM) along the primary roots of *A. thaliana* Col-0 (magenta), *A. thaliana cinv1* (teal), *S. italica* (purple), *S. viridis* (light blue) plants. Seedlings

were imaged 24 hours post transfer to Suc-MAPP-3L-embedded agar along with sucrose standard controls (0, 100, 1000 µM). **d,** Ratiometric sfGFP/mScarlet (mean ± SEM) at increasing horizontal distances from the primary root (0, 413, 828, 1655 µm) between *A. thaliana, S. italica,* and *S. viridis*.

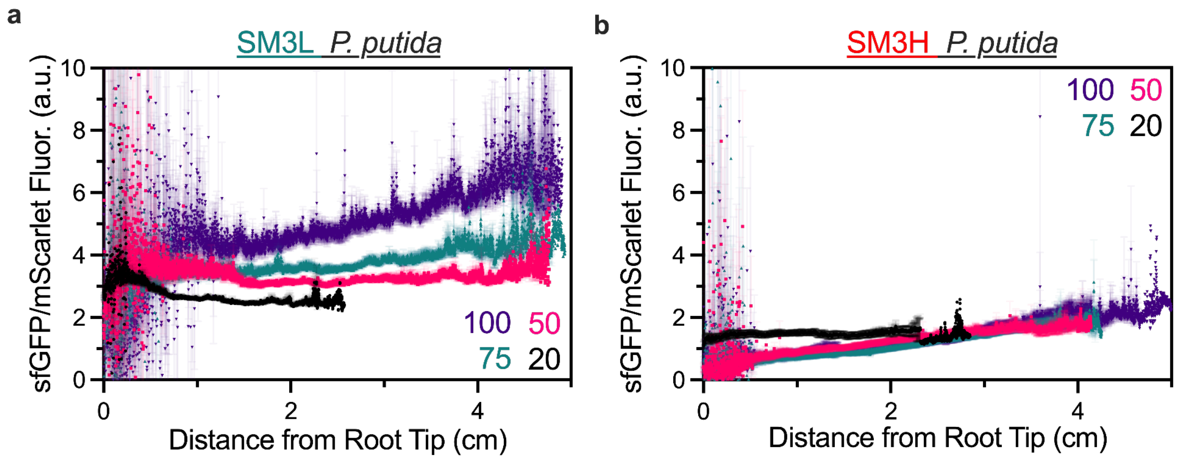


**Figure S12. *A. thaliana* sucrose exudation pattern across varying photon flux intensity.** **a,b,** Average ratiometric sfGFP/mScarlet signal ± SEM along *A. thaliana* Col-0 primary roots grown on solid plant media under varying light intensities (20, 50, 75, 100 µmol $m^2$ $s^{-1}$). Seedlings were imaged 5 days post inoculation with the *P. putida* Suc-MAPP-3L biosensor. **(b)** Seedlings inoculated with Suc-MAPP-3H biosensor.

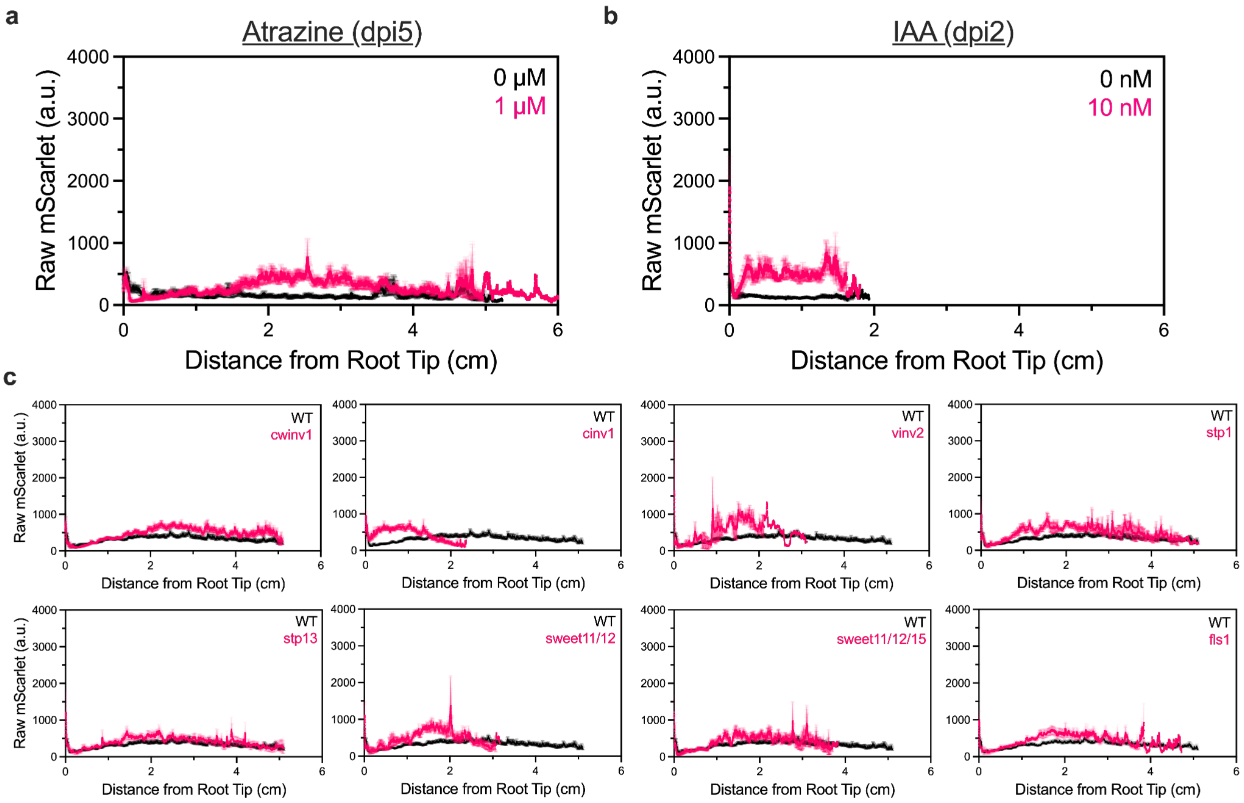


**Figure S13. Effect of sink–source perturbations to biosensor root colonization.**

**a,** Quantification of sfGFP/mScarlet fluorescence (mean ± SEM) along *A. thaliana* Col-0 primary roots (n = 12) grown on solid plant media was supplemented with atrazine (1 μM) and IAA (10 nM). All seedlings were imaged either 2 or 5 days post inoculation with the *P. putida* suc-MAPP-3L biosensor. **b,** Quantification of sfGFP/mScarlet fluorescence (mean ± SEM) along the primary roots of *A. thaliana* knockout collections referenced in Figure 3. All seedlings were imaged 5 days post inoculation with Suc-MAPP-3L.

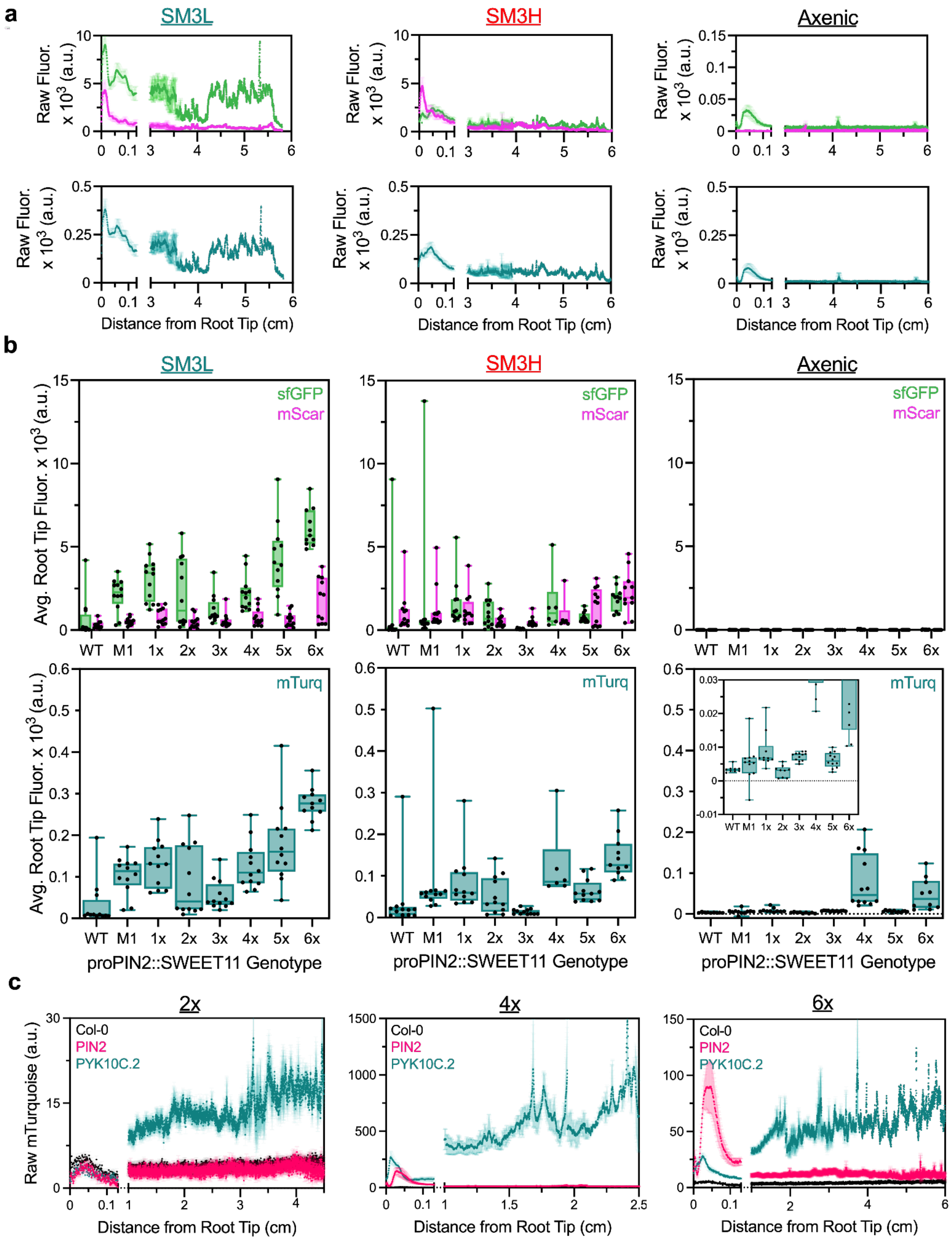


**Figure S14. Raw sfGFP, mScarlet, and mTurquoise fluorescence across axenic and biosensor-colonized Buffer Gate lines.**

**a,b,** Representative average raw sfGFP, mScarlet, and mTurquoise ± SEM signals along the primary roots of T3 proPIN2::SWEET11 6x Buffer Gate seedlings. Roots were imaged 7 days post inoculation with either *P. putida* Suc-MAPP-3L (SM3L), Suc-MAPP-3H (SM3H), or an axenic control. **(b)** Average root tip (0–1cm) sfGFP, mScarlet, and mTurquoise fluorescence (mean ± SEM) along the primary roots of all T3 proPIN2::SWEET11 Buffer Gate strengths (M1–6x). **c,** Quantification of mTurquoise2 ± SEM along the whole roots of axenic T3 proPIN2 (magenta) and proPyk10C.2 (teal) Buffer Gates (2x, 4x, and 6x) relative to a *A. thaliana* Col-0 control (black). Imaged 5 days post transfer to solid MS-sucrose medium.

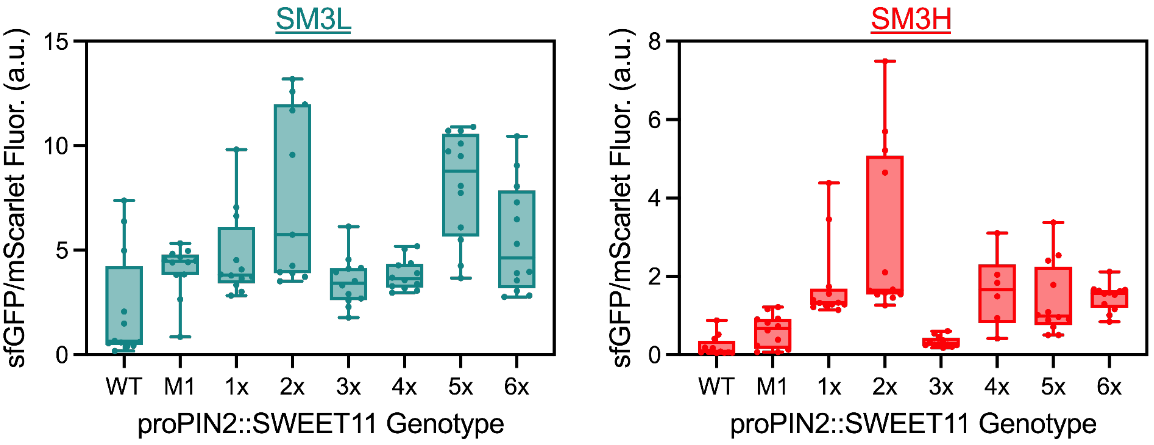


**Figure S15. T3 proPIN2 biosensor-quantified sucrose exudation**

Average root tip (0–1 cm) sfGFP/mScarlet fluorescence (mean ± SEM) along the primary roots of T3 proPIN2::SWEET11 Buffer Gates (M1, 1x, 2x, 3x ,4x, 5x, and 6x) relative to an *A. thaliana* Col-0 control (WT). Imaged 5 days post inoculation with either *P. putida* suc-MAPP-3L (SM3L) or suc-MAPP-3H (SM3H) biosensors.

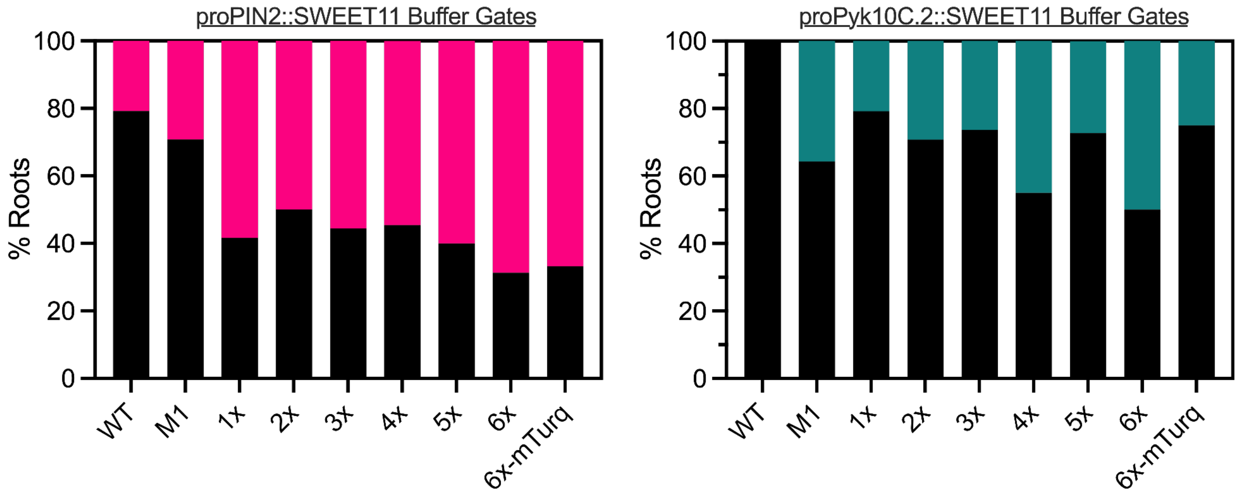


**Figure S16. Buffer Gate promoter input burden on seedling growth.**

Percentage of T1 Buffer Gate seedlings (proPIN2 or proPyk10C.2) across all strengths that did not grow 5 days post transfer to solid MS-sucrose medium. Solid black = primary root length > 1cm; solid magenta/teal = primary root length < 1cm.

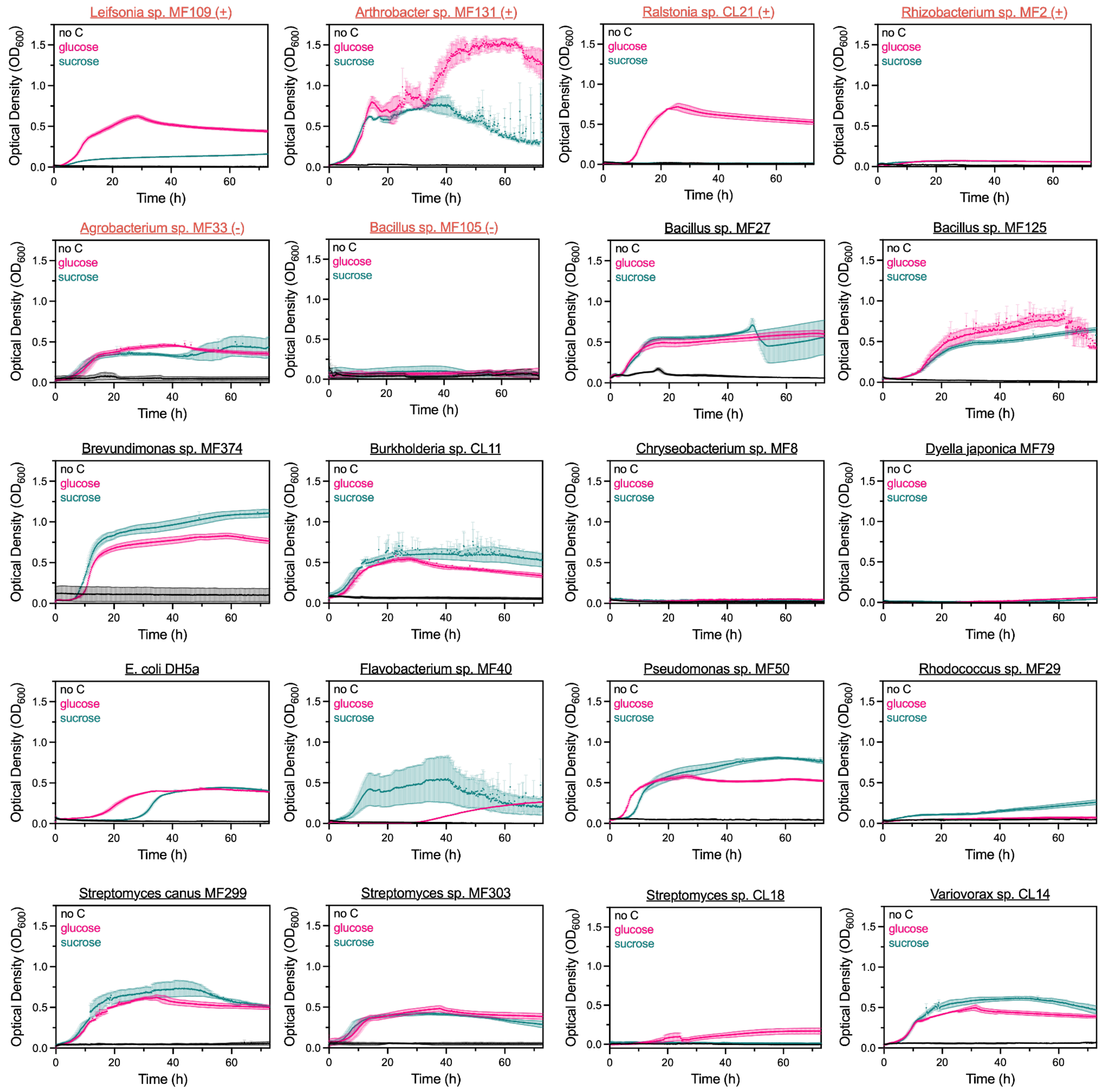


**Figure S17. Sucrose catabolism throughout the 20-member SynCom.**
SynCom strain growth, measured over a span of 72 hours, under carbon-limited conditions. Grown in liquid M9 medium supplemented with either sucrose (teal), glucose (magenta), or a carbonless control (black). Sucrose-responsive strains from Figure 5c are highlighted in orange.

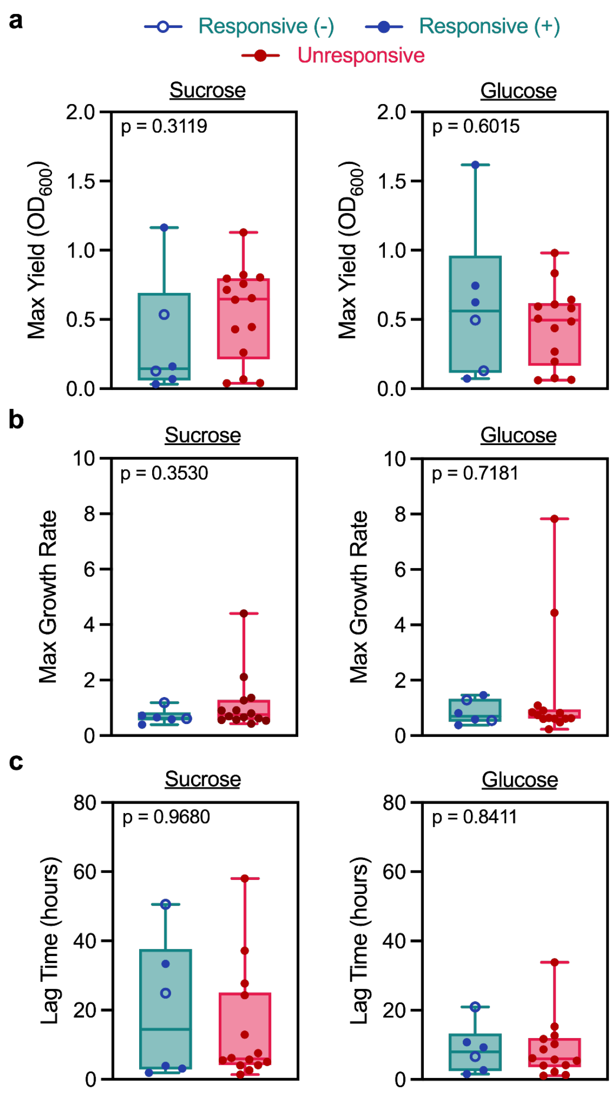


**Figure S18. SynCom carbon-limited growth trait distribution between biosensor responsiveness groups.**

Maximum yield measured as optical density ($OD_{600}$) **(a)**, maximum logarithmic growth rate **(b)**, and lag time **(c)** of SynCom strains grown in minimal liquid M9 medium supplemented with either 10 mM glucose or sucrose. Negative slope sucrose-responders (open blue circles) and positive slope responders (closed blue circles) are grouped together for comparison. Significance determined by Mann-Whitney U test ($p \leq 0.05$).

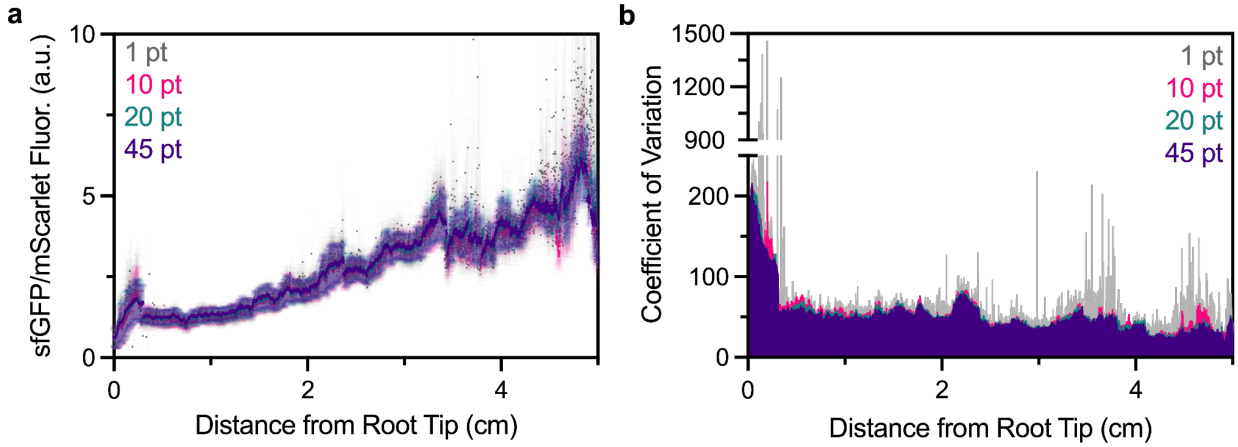


**Figure S19. Biosensor signal variation across measurement regions.**
**a,** Average sfGFP/mScarlet signal ± SEM along the primary root of *A. thaliana* Col-0 seedlings measured with different segmented line widths in Fiji: 1 pt (gray), 10 pt (magenta), 20 pt (teal), 45 pt (purple). Imaged 5 days post inoculation with the *P. putida* Suc-MAPP-3L biosensor. **b,** sfGFP/mScarlet signal coefficient of variation between n = 12 seedling replicates for each segmented line width measurement along the primary root.

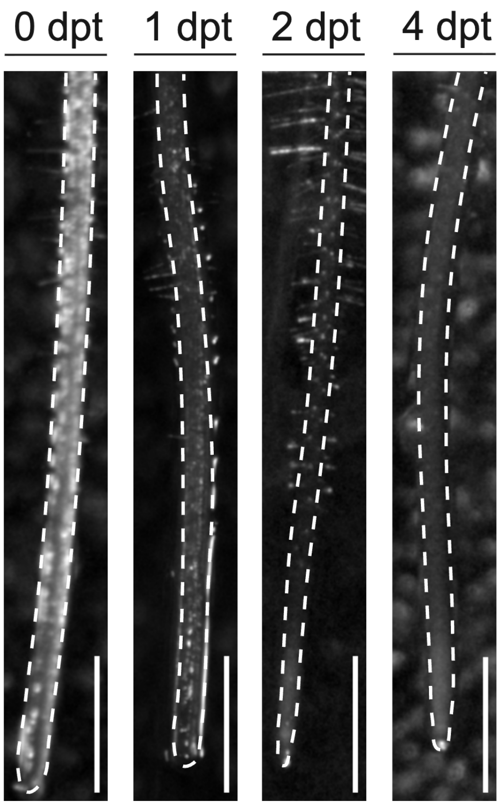


**Figure S20. Brightfield lac biosensor root outline.**
Dotted primary root outline used for Figure 2d fluorescence ratio images. Scale bar = 1 cm.

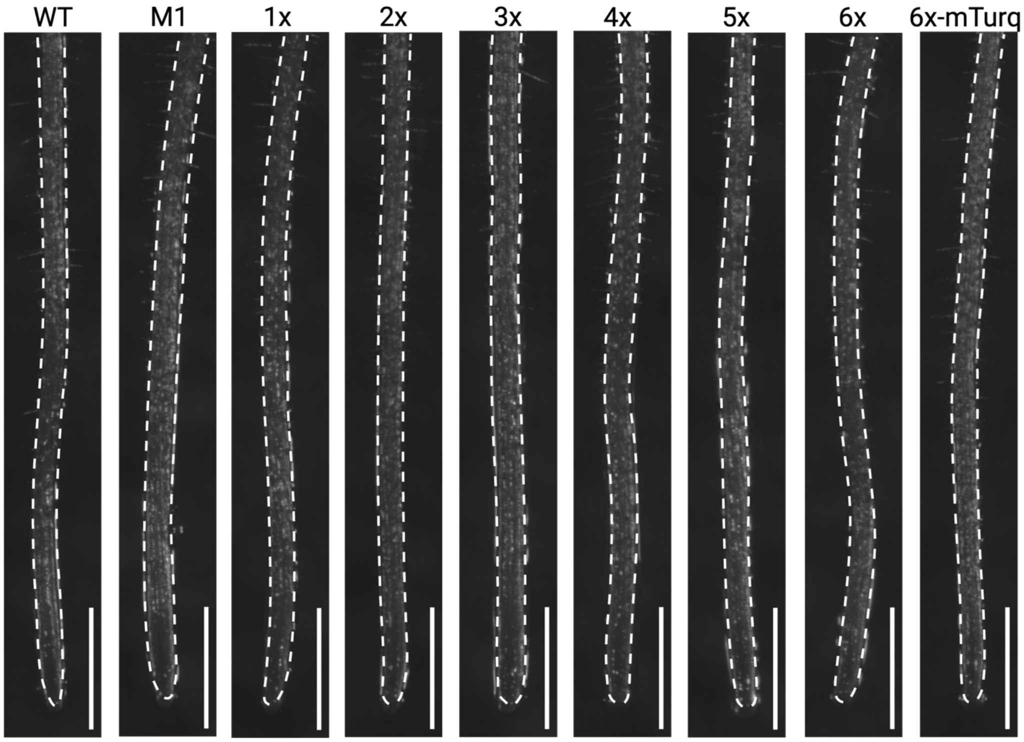


**Figure S21. Brightfield T1 proPIN2 root outline.**
Dotted primary root outline used for Figure 4f fluorescence ratio overlay images. Scale bar = 1 cm.

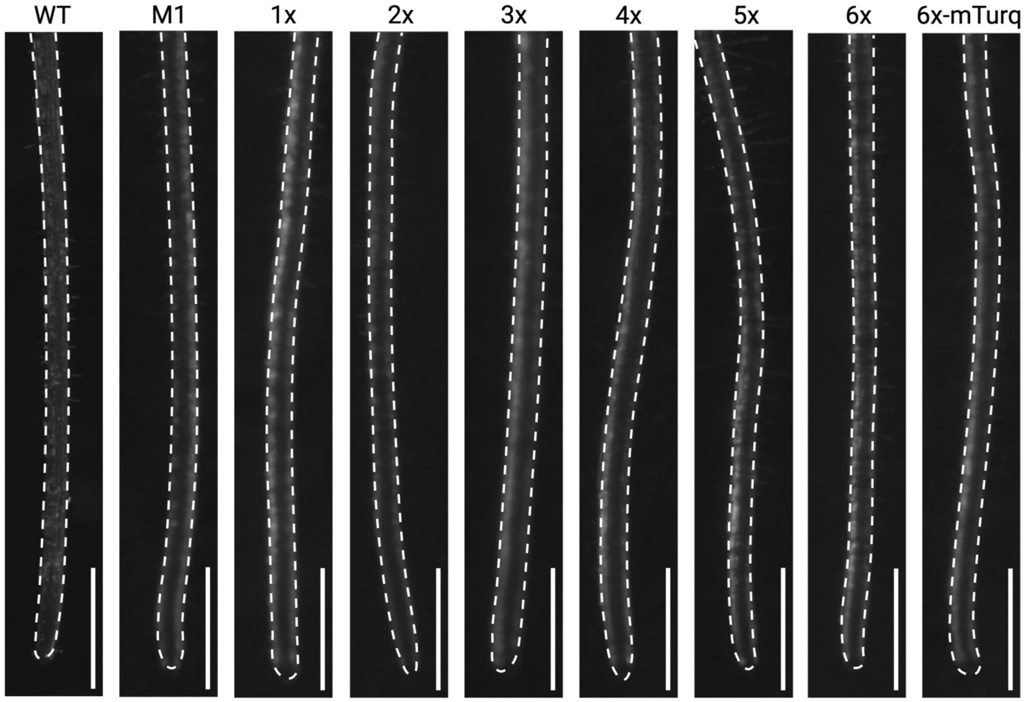


**Figure S22. Brightfield T1 proPyk10C.2 root outline.**
Dotted primary root outline used for Figure 4g fluorescence ratio overlay images. Scale bar = 1 cm.

| Taxon ID | Genus | Species | Strain | Putative Sucrose Transporters | Putative Sucrose Invertase | Sucrose Metabolism Confidence |
|---|---|---|---|---|---|---|
| 1279031 | *Agrobacterium* | sp. | MF33 | EYD00_RS01180 (aglE), EYD00_RS01185 (aglF), EYD00_RS01190 (aglG), EYD00_RS01200 (aglK), EYD00_RS21390 (thuF), EYD00_RS21080 (thuG), EYD00_RS21075 (thuK) | EYD00_RS01195 (ams) | High |
| 1157944 | *Arthrobacter* | sp. | MF131 | C566_RS0118520 (aglK), C566_RS0119160 (thuF), C566_RS0119155 (thuG), C566_RS0118520 (thuK) | C566_RS0117125 (ams) | High |
| 1404 | *Priestia* | *megaterium* | MF27 | H543_RS0100295 (cscB) | H543_RS0100290 (ams) | High |
| 1150400 | *Brevundimonas* | sp. | MF374 | BLR19_RS00805 (aglK), BLR19_RS00805 (thuK) | BLR19_RS06185 (ams) | Medium |
| 1304868 | *Flavobacterium* | sp. | MF40 | -- | -- | -- |
| 1150399 | *Leifsonia* | sp. | MF109 | H347_RS0104495 (thuF) | H347_RS0109735 (ams) | Medium |
| 1380362 | *Ralstonia* | sp. | CL21 | ABZR87_RS18670 (aglK), ABZR87_RS18670 (thuK) | ABZR87_RS17220 (ams) | Low |
| 668369 | *Escherichia* | *coli* | DH5a | NJ74_RS00950 (aglK), NJ74_RS00950 (thuK) | NJ74_RS10875 (ams) | Low |
| 32008 | *Burkholderia* | sp. | CL11 | -- | -- | -- |
| 1172181 | *Streptomyces* | sp. | MF303 | H294_RS0103425 (thuG) | H294_RS0110160 (ams) | High |
| 1172182 | *Streptomyces* | sp. | MF136 | BLQ64_RS12750 (aglK), BLQ64_RS24540 (thuG), BLQ64_RS12750 (thuK) | BLQ64_RS17870 (ams) | Low |
| 198618 | *Pseudomonas* | *umsongensis* | MF50 | AF70_RS03890 (thuE), AF70_RS03885 (thuF), AF70_RS03880 (thuG), AF70_RS03875 (thuK) | AF70_RS03870 (ams) | High |
| 1172183 | *Streptomyces* | *canus* | MF299 | none | H293_RS0125525 (ams) | High |
| 1449077 | *Streptomyces* | sp. | CL18 | BR33_RS0128615 (thuG) | BR33_RS0120285 (ams) | Medium |

| | | | | | | |
|---|---|---|---|---|---|---|
| 1246459 | *Rhizobacterium* | sp. | MF2 | D460_RS0105365 (thuE), D460_RS0105360 (thuF), D460_RS0105355 (thuG), D460_RS0105350 (thuK), D460_RS0110305 (aglE), D460_RS0110310 (aglF), D460_RS0110315 (aglG), D460_RS0110325 (aglK) | D460_RS0110320 (ams) | High |
| 1340435 | *Chryseobacterium* | sp. | MF8 | none | none | Low |
| 1380371 | *Bacillus* | sp. | MF125 | N521_RS18420 (aglK), N521_RS18420 (thuK) | N521_RS03915 (ams) | High |
| 1380365 | *Dyella* | *japonica* | MF79 | N515_RS0114025 (aglK), N515_RS0114025 (thuK) | N515_RS0111055 (ams) | Medium |
| 1151120 | *Bacillus* | sp. | MF105 | C564_RS0106745 (aglK), C564_RS0118840 (thuK) | C564_RS0120250 (ams) | High |
| 1150676 | *Rhodococcus* | sp. | MF29 | -- | -- | -- |
| 1150395 | *Variovorax* | *paradoxus* | CL14 | D461_RS0106115 (aglK), D461_RS0119620 (thuG), D461_RS0106115 (thuK) | D461_RS0109270 (ams) | High |

**Table S1. Gap-Mind Predicted Sucrose-Utilizing Taxa.** Analysis of SynCom strains with GapMind[1] to predict sucrose uptake and catabolism. High-confidence predictions (green), medium-confidence predictions (orange), low-confidence predictions (magenta).

**Bacterial Strains**

| GENUS | SPECIES | STRAIN | PLASMID;GENOTYPE | SOURCE |
|---|---|---|---|---|
| *Pseudomonas* | *putida* | KT2440 | n/a;WT | Andrew Ellington |
| *Pseudomonas* | *putida* | Empty Plasmid | pCDrRK3; $P_{ECrrnB(disc)}$::*mScarlet-I* | This manuscript |
| *Pseudomonas* | *putida* | $P_{sux2}$::*sfgfp* | pCDrRK70; $P_{ECrrnB(disc)}$::*mScarlet-I*, $P_{sux2}$::*sfgfp* | This manuscript |
| *Pseudomonas* | *putida* | Suc-MAPP-1 | dCMD24; $P_{ECrrnB(disc)}$::*mScarlet-I*, $P_{J23107}$::*suxRcscBY*, $P_{sux2}$::*sfgfp* | This manuscript |
| *Pseudomonas* | *putida* | Suc-MAPP-2 | dCMD30; $P_{ECrrnB(disc)}$::*mScarlet-I*, $P_{J23107}$::*suxRcscBY*, $P_{sux1}$::*sfgfp* | This manuscript |

| | | | | |
|---|---|---|---|---|
| *Pseudomonas* | *putida* | Suc-MAPP-3L | dCMD101; *$P_{ECrrnB(disc)}$::mScarlet-I, $P_{J23107}$::suxRAC, $P_{sux1}$::sfgfp* | This manuscript |
| *Pseudomonas* | *putida* | Suc-MAPP-3H | dCMD102; *$P_{ECrrnB(disc)}$::mScarlet-I, $P_{J23107}$::cscRsuxAC, $P_{csc1}$::sfgfp* | This manuscript |
| *Pseudomonas* | *putida* | *lac* Biosensor | dCMD204; *$P_{ECrrnB(disc)}$::mScarlet-I, $P_{J23107}$::lacI, $P_{lac}$::sfgfp* | This manuscript |
| *Pseudomonas* | *putida* | $P_{lac}$::*sfgfp* | dCMD207; *$P_{ECrrnB(disc)}$::mScarlet-I, $P_{lac}$::sfgfp* | This manuscript |

**Plant Lines**

| **GENUS** | **SPECIES** | **Accession** | **GENOTYPE** | **SOURCE** |
|---|---|---|---|---|
| *Arabidopsis* | *thaliana* | Col-0 | WT | Dinneny Lab |
| *Arabidopsis* | *thaliana* | SALK_095807 | *cinv1* | ABRC |
| *Arabidopsis* | *thaliana* | CS25124 | *cwinv1* | ABRC |
| *Arabidopsis* | *thaliana* | SALK_100813 | *vinv2* | ABRC |
| *Arabidopsis* | *thaliana* | SALK_141277 | *fls2* | Libo Shan |
| *Arabidopsis* | *thaliana* | SALK_048848 | *stp1* | ABRC |
| *Arabidopsis* | *thaliana* | SALK_021204 | *stp13* | ABRC |
| *Arabidopsis* | *thaliana* | | *sweet11/sweet12* | Wolf B. Frommer |
| *Arabidopsis* | *thaliana* | | *sweet11/sweet12/sweet15* | Wolf B. Frommer |
| *Arabidopsis* | *thaliana* | PIN2–M1 | proPIN2 Buffer:: SWEET11–*mTurq2* (M1) | This manuscript |
| *Arabidopsis* | *thaliana* | PIN2–1x | proPIN2 Buffer:: *SWEET11–mTurq2* (1x) | This manuscript |
| *Arabidopsis* | *thaliana* | PIN2–2x | proPIN2 Buffer:: *SWEET11–mTurq2* (2x) | This manuscript |
| *Arabidopsis* | *thaliana* | PIN2–3x | proPIN2 Buffer:: *SWEET11–mTurq2* (3x) | This manuscript |
| *Arabidopsis* | *thaliana* | PIN2–4x | proPIN2 Buffer:: *SWEET11–mTurq2* (4x) | This manuscript |
| *Arabidopsis* | *thaliana* | PIN2–5x | proPIN2 Buffer:: *SWEET11–mTurq2* (5x) | This manuscript |

| | | | | |
|---|---|---|---|---|
| *Arabidopsis* | *thaliana* | PIN2–6x | proPIN2 Buffer:: *SWEET11–mTurq2* (6x) | This manuscript |
| *Arabidopsis* | *thaliana* | PIN2–6xmT | proPIN2 Buffer:: *mTurq2* (6x-mTurq) | This manuscript |
| *Arabidopsis* | *thaliana* | PYK–M1 | proPYK10C.2 Buffer::*SWEET11–mTurq2* (M1) | This manuscript |
| *Arabidopsis* | *thaliana* | PYK–1x | proPYK10C.2 Buffer::*SWEET11–mTurq2* (1x) | This manuscript |
| *Arabidopsis* | *thaliana* | PYK–2x | proPYK10C.2 Buffer::*SWEET11–mTurq2* (2x) | This manuscript |
| *Arabidopsis* | *thaliana* | PYK–3x | proPYK10C.2 Buffer::*SWEET11–mTurq2* (3x) | This manuscript |
| *Arabidopsis* | *thaliana* | PYK–4x | proPYK10C.2 Buffer::*SWEET11–mTurq2* (4x) | This manuscript |
| *Arabidopsis* | *thaliana* | PYK–5x | proPYK10C.2 Buffer::*SWEET11–mTurq2*(5x) | This manuscript |
| *Arabidopsis* | *thaliana* | PYK–6x | proPYK10C.2 Buffer::*SWEET11–mTurq2* (6x) | This manuscript |
| *Arabidopsis* | *thaliana* | PYK–6xmT | proPYK10C.2 Buffer::*mTurq2* (6x) | This manuscript |
| *Brassica* | *rapa* | PI 633171 | | NPGS |
| *Brassica* | *napus* | Darmor-bzh | | Dinneny Lab |
| *Setaria* | *italica* | PI 690671 | | NPGS |
| *Setaria* | *viridis* | WV001-9 | | NPGS |
| *Solanum* | *pennellii* | PI 365970 | | NPGS |
| *Sorghum* | *bicolor* | *BTx623* | | *Katrien Devos* |

**DNA**

| **PART TYPE** | **PART NAME** | **DNA SEQUENCE** |
|---|---|---|
| Bacterial Promoter | $P_{sux1}$ | TAGGTGTTTACAGCTAGCTCAGTCCTAGGTATTATCATTGCAATCGATTGCATTG |
| | $P_{sux2}$ | CTCGTTGCAATCGATTGTAACGTTGACATTGCAATCGATTGCATTGATACTGAGCAC |

| | | |
|---|---|---|
| | $P_{csc1}$ | TAGGTGTTTACAGCTAGCTCAGTCCTAGGTATTATCGATGTTA<br>ACGTTAACCTAA |
| | $P_{J23107}$ | TAGGTGTTTACGGCTAGCTCAGCCCTAGGTATTATGCTAGCT<br>CTAGA |
| | $P_{ECrrnB(disc)}$ | TAGAGTTGACTAAGACTTGTCAGGCCGGAATAACTCCCTATA<br>ATGATACACCA |
| | $P_{lac}$ | TAGGTGTTTACAGCTAGCTCAGTCCTAGGTATTATGAATTGT<br>GAGCGCTCACAATTC |
| Bacterial Gene | *suxR* | ATGGCTAAACATCCTGAAAGGGCCACGCCCGCGGCGCGACC<br>ACAAATGGCGGATATTGCTCGCATGGCTGGCGTATCGGAGT<br>CAACGGTATCGCGAGCGCTTGCAGGCTCGCCTGTAGTCGCG<br>GAGCGCACCAGGGCCTACATCAAACAGATCGCCGCTGACGC<br>CGGCTACCAAGTCGATCCTGTAGCTCGCAGCCTTCGCGCGA<br>AGCGATCCAATACAGTGTCAGTCGCTGTTCCCATGATGCATG<br>CTTTGGATCAACCGCTTTCGGATCCGTTTCTGATGACGATGC<br>TTGCCTTGCTGGCGGAAGAGTTGACAGGTCGGGGTTACTCA<br>ATGCTTCTCAGCAAACTGAACCGACATCAGGACGGTTGGGTT<br>GAACAGTTGGCTCGCGGATCACGGAGCGATGGCGTCATCGT<br>CTTGGGGCAATCCTCCGAACACGCCGCCTTGGACGCCGCTG<br>CGCGGGATGGCCTCCCCATGGCCGTTTGGGGCTCGCGTATC<br>GACGGCCAGTCCTACATATCGGTCGGTAGCGACAATTTCCA<br>GGGCGGGGCTCTCGCTACTGAGCACTTGATCGCTAGCGGCC<br>GGCGGCGCATCGCCTTTCTCGGGGACGATCAATTGCCGGAA<br>GTCGCCCCGCGGTTCGCGGGATACAGGCAGGCCTTGGAGC<br>AGCACGGGTTGGAATTCGACACCAGGCTGCATGCTCGGTCA<br>CATTTTCTTTCCGAGGATGCTTATCGACTCACAAGGGCCATG<br>TTGAAAAAGACCGACGCCCCAGACGGCTTGTTTGCCGCTTC<br>AGACGTGATCGCAGTAGGTGCGATTCGTGCCCTCGTCGAGG<br>CGGGCCACCGCGTTCCACAAGATATCTCGTTGGTCGGTTTC<br>GACGACATTCCTTTGGCTGCGTATAGCCAACCACCTCTTACC<br>ACCGTTCAGCAAGACCTCGGACTCGCGGCACGACTCCTGGT<br>GGACCGATTGCTCGCGCTCATCGCCGGTGAGGCCGTTGACA<br>GCGTTGAGATGCCCGTGAAACTCGTGGTCCGGGAATCGGCT<br>TAA |
| | *suxA* | ATGTCCACCTTGCACACCCTGCGCCTGCACGCTTTGGCCTG<br>CGCGGTCACCACCTGCCTGTGCGCGCCGCTCGCGCTGGCC<br>CAGGACGCCACCCCGCCCGCACCCGCCACCCCGCCGGCCG<br>CCGACAGCGCCGCGGTCAACCTGGATTCGGTGTTCGTCACC<br>GGCACCTCCACCGCCACCACCAAGCTCAAGTCCAGCGTGTC<br>GGTCAGCACCGTCGGTGCCGAGGCGATCGAGCAATCGGCC<br>CCGCGCAGCACTGCAGAAATCTTCCGCAACATTCCCGGTAT<br>CCGCTCCGAGTCCAGCGGCGGCGAAGGCAATGCCAATATCG<br>CCGTGCGCGGCCTCCCCGTCGCTTCCGGCGGCGCCAAGTT<br>CCTGCAGCTGCAGGAAGACGGCCTGCCGGTGATGGAGTTC<br>GGCGATATCGCCTTCGGCAATGCCGATATCTTCCTGCGCTC<br>GGATTTCACGATGGATCGCATCGAAGCGATCCGCGGCGGTT<br>CGGCCTCCACGTTTACCAGCAATGCGCCCGGCGGCATCATC<br>AACTTCATCAGCAAGACCGGCGACACCGAAGGCGGCAGCGT<br>CGGCGTCAGTCGCGGCCTGGACTACGACAACACCCGTATCG<br>ACTTCAATTACGGCGCGCCGTTTGCCGAGCACTGGCAGTTC |

| | | |
|---|---|---|
| | | AACATCGGCGGCTTCTTCCGCCAGGGCGACGGTGTGCGCG<br>ATGCCGGCTACACCACCGACAAGGGCGGCCAACTCAAGGCC<br>AACCTCACCCGCCTGTTCGAGAACGGCTACGTGCGTGTGTA<br>CGGCAAATACCTCAACGACCGTGCCGCCGGCTATTTGCCGG<br>TGCCCACCTCGGTGCGTGGCCGCGACGGCTCGCCGGACCT<br>GGGCGGCTTCCCCGGTTTCGACCCCGGCAACGACACCTTGT<br>ACAGCCGTAATTTCCGCACCGATGTTGGCCTGGATGGCAAC<br>AACCAGCCGCGCCGCACCGACCTGGGCGACGGCATGCACC<br>CGATCTCCCGCACCATCGGCGCCGAAGCCTGGTTCGACCTT<br>GGCAATGGCTGGAACCTGAGCGATAAATTCCGCATCGCCGA<br>CAACAGCGGCCGCTTCGTCAGCCCGTTCCCGGCCGAAGTCA<br>CCGATGCCGCCGCGCTGGCATCGTCCATCGGCGGCGCCGG<br>CGCGCAGCTGGTGGAAGCCGGTGGCGCCAATGCCGGCCAG<br>GCCTACACCGGCACCGCCATCCGCACGCATCTGTTCAATGT<br>TGCCATCAACGATCTGGGCAATGTCACCAACGACCTCAGCCT<br>CTCGCGCGAGTTCGGTGGTGACGGCCGCACCCTCAACCTGC<br>GCATGGGCTACTACACCTCGCGCCAGACCATCGACATGGAC<br>TGGACCTGGAACTCCTACGTGCAGAGCCTGGGCCGCGACTC<br>GCGCCTGCTCAACGTGGTCGACGCCAGCGGTGTGTCGCGC<br>TCGCAGAACGGCCTGTACGCCTACGGCACGCCGTTCTGGGG<br>CGATTGCTGCATCACCCGCAGCTACGACGTGCGCTACGACG<br>TCAACGCACCGTACGTGGCGCTCACCTTCGACAGCGGCAAG<br>CTCAGCATCGACGGCAGCCTGCGCTACGACATGGGCGATGC<br>GCGCGGCAACTATTCCGGCACCGCCATCGCGCAGAACCTGG<br>ACGTCAACGGCGATGGCGTGATCCAGCCGGTGGAGCAGCG<br>CGTGGCCACCGTGGACACCGCCAATGCGCGGCCAGTGGAC<br>TACGACTGGAACTACCTGTCGTACTCATTGGGCGGCAATTAC<br>CTGATCAACGACGACCTGGGCGCATTCGCACGCGTCAGCCG<br>TGGCGCACGCGCCAATGCCGACCGCCTGCTGTTCGGCGTG<br>ATCCGCGACGATGGCTCGGTGACCTCCGATGAAGCGGTCAA<br>CGTGGTGCGACAGACCGAAGCCGGCCTGAAGTGGCGCCGC<br>GACGGCCTGAGCCTGTTTGCCACCGCGTTTGCCGCGCGCAC<br>CCAGGAGCAGAACTTCGAAGTCACCAGCCAGCGCTTCTTCA<br>ACCGTAGCTACAAGGCGCACGGGATCGAGCTGGAAGCCAG<br>CTACCGCTACGAGGGCTTCACCGTCAACGGCGGCGTGACCT<br>GGACCGATGCAGAAATCGCCAGGGACCAGATCACTCCGGAG<br>AACACCGGCAACGTGCCGCGCCGCCAAGCCGACTTCGTGTG<br>GCAGCTCACCCCGAGCTACCGCGGCGACGGCTACCAGTTC<br>GGCGTGAACCTGATCGGCACCACCGAGGCCTACACGCAGG<br>ATTCCAACCAGCTGAAGATGCCGGGCTACACGCAGGTGAAC<br>CTGTTCGGCGACTACCGCATCACCGATGCGCTCACCGTCGC<br>GTTGAACGTCAACAACCTGTTCAATACGTTCGGGCTGACCGA<br>AGCCGAGGAAGCCACCATTCCGGCCAACGGCATCATCCGCG<br>CACGTTCGATCGCCGGCCGCACCACCAGCCTCAGCCTGCGC<br>TACGACTTCTAA |
| | *suxC* | ATGCCTGGACCCTGCATGTCGTCGACCGCTCCTCCGCTTTC<br>CTTCGCACGCATCCTGGCCCTCAATGCCGGGTTCTTCGGTG<br>TGCAATACAGCTTCGGGCTGCAACAGAGCAACATGAGCCCG<br>ATCTACAACTACCTGGGTGCCGACCACGCCAGCCTGCCGTA<br>CCTGTGGCTGGCCGGGCCGATCACCGGCCTGGTGCTGCAG<br>CCGTTCGTCGGCGCCTTGAGCGATCGCTCGGTGACGCGCTG<br>GGGCCGGCGCATGCCCTACATGGTGCTCGGCGCGTTGGTG<br>TGCAGCCTGTGCCTGCTGGCGATGCCTTTCAGTACCGCATT<br>GTGGATGGCGGTCTGCCTGCTGTGGGTGCTGGACGCCGCA |

| | | |
|---|---|---|
| | | AACAACGTCGCGATGGAGCCGTACCGCGCCCTGGTCAGCGA<br>CGTGCTGGCGCCACCGCAGCGGCCGTTGGGGTATCTCACC<br>CAGAGCGCCTTCACCGGGCTGGCGCAGACGCTGGCCTATCT<br>CACCCCGCCGTTGCTGGTGTGGTTCGGCATGAGCCAGGACG<br>CGGCCAATGCGCACCACATCCCGTACGTGACCATTGCTGCG<br>TTCGTGATCGGCGCCGGCTTTTCGGCCGCCTCGATCCTGCT<br>CACCGCGCGCAGCGTGCGCGAGCCGGTGGTGCCTGCGGCG<br>GAGATCGCGCGCATGCGCAAGGCCGGTACCGGGCTGGGCG<br>CCACGCTGCGCGAGATCGGCAGTGCGCTGCGCGACATGCC<br>ACCGACCATGCGCCAGCTGGCGCCGGTGATGCTGTTTCAGT<br>GGTATGCGATCTTCAGCTACTGGCAGTACATCGTGCTGTCGC<br>TGTCGACCACCTTGTTCGGCACCACCGAGGCGAACTCGCAC<br>GGCTTCCGCGAGGCCGGGCTGGTCAACGGGCAGATCGGTG<br>GTTTCTACAATTTCATCGCGTTTCTGGCGGCTTTTGCGATGG<br>TGCCGGTGGTGCGGCGCGTCGGCCCCAAGTACACGCACGC<br>GGCCTGCCTGCTGGCGGCTGGCGTGGGCATGTGGGTGTTG<br>CCGGGCATCGAGAACCGTTGGTTGTTGCTGCTGCCGATGAT<br>CGGTATTGGCCTGGCCTGGGCCAGCATGATGGGCAACCCCT<br>ATCTGATGCTGGCCGATAGCATTCCGCCCGAGCGCACCGGC<br>GTGTACATGGGCCTGTTCAACCTGTTCATCGTGCTGCCGATG<br>CTGATCCAGATCGTGACCTTGCCGCTGTATTACGAACCGCTG<br>TTGCACGGTGACCCGCGCAATGTGATCCGCCTGGCCGGCG<br>CACTGATGCTAGCGGCTGCGGTGGCCATGCTGTGCGTACGC<br>ATCCGCAAACCCAAGGCCGTGGCGGCCTAG |
| | *cscR* | ATGGCCTCAGTTAAAGATGTCGCGCGCCTGGCAGGGGTTTC<br>CTTCATGACCGTGTCCAGGGCGCTTAACACTCCGGAAAAGG<br>TCAATCAGGAAACCCTGGCGAAAGTTCTGCAAGCGGTTGAAA<br>CCCTGGGTTATGTTCCCAGCCTTTCTGCCCGAAAGATCCGC<br>GGCGGTGCTTCCCGTGGCAAGACCATTGGCGTGTTCGCGCT<br>GGATACCGCCACCACGCCCTTCGCTGTGGAAATGCTCCTGT<br>CCATGGAGCGCACTGCCCGGGAGCACGGCTGGAATATCTTC<br>ATCCTCAACGTGTTCGAAGTGCCGCCGAGCCAGCAGACCAT<br>CGACCTGATGCTGTCCCACCAGCCTGACGGAATCATTTTTAG<br>CGCCATGCAGTTGCGGACTGTCGAAATCCCCGCAGGGCTGC<br>GCAGCCTGCCGTTGGTACTGAGCAATTGCGTGAGCCTGGAG<br>CCCGGGGTGGCCTGCTATGTGCCTGACGATGCCGATGGCCA<br>GTATCAGGCGGTGCGACATGCGCTCAAGCGTGGTTATCGGC<br>GTCCGTTGTGCATCAACCTGCCACAAGCCAGCCTGGCTTGG<br>CAGCCGCGCCAGCTTGGATTGTGTCGCGCACTCGCCGAGG<br>CCGGAATATCCGTGGCCGATGTTCCTCAATACAGCCTGTCG<br>CAGGATGACGCCTATCAGGAAACGCTCAGCGTGCTGGAGAC<br>TCAGTTGCGCGAGTCGCAAGAAGGTCCAGCGTTCGATCTCC<br>TTATCTGTGGTAACGACCGTATTGCTTTGGTGGCCTACCAGT<br>ACTTGCTAAGCCGTGGGCTGCGTATTCCGAACCAGGTGGCG<br>GTCTTGGGTTACGACAACATGATCGGTGTTGCAGAGCTGTTC<br>TACCCGCCGCTCAGTACTGTGCAGTTGCCGTATTACGAAATG<br>GGACGACGCTCTGTCCAATACATCATTGAGTGCAGGAACGA<br>ACCCGGGATTCAACGGGTTGAATGCCCCGTAGTTGAACGTG<br>AGTCGTGTTAG |
| | *cscB* | ATGCAGTTTGCAGCCAAACGGGAGTACTGGTTGATCAGCGG<br>GCTTTTGTTCTTCTTTTTCTTTTCCTGGAGCTCCTCCTACAGC<br>TTGTTCAGCATCTGGCTTCACCGTGTCATTGGACTCAATGGA<br>ACAGAGACAGGCTTCATATTCGCAGCAAATGCAATCGCTGCT |

CTGCTCGTACAACCATTCTACGGAGCACTGCAAGATCGGTTG
GGGCTCTCCAAGAAACTTTTGGTGTGGATTGGGATACTCCTT
TGCGCCGCTGCACCCTTCGCTATCTATGTATACGCGGGGCT
TTTGGCACAGAATGTTATGCTCGGAGCATTGGTCGGGGCTG
CCTTCTTGGCTTTGGCTATGCTGGCAGGAGTGGGAGTGATT
GAGAGCTATACCGAGAGGCTCAGCAGGCACGCCGGTTTTGA
ATTCGGGACTACACGTATGTGGGGCAGCTTGGGTTGGGCTA
GCGCGACAGGCGTTGTTGGAGTGGTCTTTAACATTGATCCTG
ACATTGCCTTCTATATGTCGTCGTTGGCAGGAATCGTATTCC
TTTTGATCCTGTTCCGCTTGGATCTCGACCGTCTGGCTCAGC
CCGCAGTCCAAGCAGGAGCTGTGGTTCACCCTGTCCGACTT
AACGACCTTTGGAAACTGCTGGCCCTTCCACGGTTTTGGGCT
TTTAGCTTGTACCTCACTGGAGTATGTGGAATCTACATGATCT
ATGAACAGCAGTTTCCCGTCTACTTTTCATCGTTTTTCCCCAC
GCCTGAGGAAGGAACGCGTGCGTACGGATATCTCAACTCGT
CCCAAGTATTGGTGGAAGCGGTACTTATGTTGTTGGCCCCGT
GGGTTGTTTCCCGAACAGGTGCGAAATACGGCCTCATATTG
GCTGGTAGCATCATGTTCGTCCGAATTTTGGGGTCGGGTCTT
GTTACTCAAGCGTGGGCGATAGCTGCTTGTAAGATGTTGCAT
GCTCTTGAGGTACCGATCCTGCTTGTCTCGATTTTCAAATAC
ATATCCCTGAATTTCGACAGCCGGCTTAGCGCGTCGATTTAC
CTTGTAGGATTTCAATTTGCCCAACAGCTGACTGCGATGTTG
CTGTCGCCCCTTGTAGGATACGGATACGACCACTTTGGGTTC
TCGTCAGTGTACGTTCTTATGGCTGGCTTGGTCGGGGCGTG
TCTCTTGTTGTCGTGGACACTGCTTCGTAAGGACCCAGTTCG
CGATGCCTCGCAAGTGGGTGCTGGGGACTCCAGGCAGTTGC
CTGCAATCGCTCCTTCAGCCCCTCGTTACGAACCTTAG

| | |
|---|---|
| *cscY* | ATGCGCAGCGCGGTTTGCTGGCCATGGGCTGGATTCAGCTT<br>GTTTGTTCTTTCGACACAATTGTTTGCGGCCCCGGCACAAAG<br>CATTGAAGAACGCCTTGCAAGGCTGGAAGCTCGAACCTCGA<br>GCGCGGAAGCCCGAGCGAGCGCTGCCGAGGCGGATGCCG<br>CACGCCTGCGTCGCGAGGTCCAACTGCTCAATCAAAAGATC<br>GTGGGCCGTTTGCCCAGCCCAGCAGAGAATGCTCTGGACCA<br>GCGCATAGCCAGGATTGAAGCCCATCAACAAGTGCTCGCGT<br>CCGCCGCACCCGCTACAGCATCATCAGCCGGCGCGAGCGA<br>ACGGCTTTCGGACGGTTTCACTTTTGGTGGTTATGCCCGCAG<br>CGGCATCATGTCGAATGGCTCCGGAGCAGGACGGGGTGGA<br>CCATATGTTACCCCGGCTGGGAGCGTGGGTGGTGCAGTTGG<br>TAGGTTGGGTAACGAAGTCGACACCTACATGGAAGCGAAGC<br>TCGCTAAGGAAAGCCAGGCAGATAACGGCACTCATGCTAAG<br>TACTTGTTGATGCTGGCGGACGGTCTGGAAACACCTAACGA<br>CTGGACAGCGGCACAATCCCAACTGAATGTGCGACAAGCAT<br>ACACTGAGTTGAGCCACTTGGCGTCGTTTCAAGATAGCCCAC<br>TGCTTCATAACGCGACTTTGTGGGCAGGGAAACGCTTTGACC<br>GTGACAACTACGATCTTCATTGGCTGGACCGTGGGATCGTCT<br>TTCTGGCCGGTACTGGTGGCGGAATATATGACCTGCAGTTG<br>ACGCAAGACTGGCGACTTAATGCCAGCTTGATGTCGCGCTC<br>CTATGGTGATTTTGGAACAGAGGAAAAGAAAGACATACGCAG<br>CTACGTGGCGACGCTTAATCAGTTCTTCGATCAGGGACGGT<br>GGCAGGTGATGTTGAATGGGATTTCATCAGGACAGAACGAC<br>GCTGACTTGGAACCGGGGAACGGGGCAGAACCGGACCGGA<br>AGCGGCGGCTGAACAAGTCCGGGTTTTCGCCGGCCACCGG<br>TGGCACACACGGGCTTCTGGCATATCACCGTCCTGACTTCTT<br>TGGCCATGAGGGTTACGCGAAGGCTGTTTTGTTGTATGGACA |

AGGCTTGGGTGCAGAGGTCAACAACATTGGCGCAGATGGCG
ACCTGCTCGACCAAGCGCGGACACTTCGTCTGGCGTTCTAT
GGACACACGCGATTGAATCAGGACTGGCGGATCGCACCGGC
ACTTATAGCCGAACAAAGCAAAGATCGCTATGTCGCGGGTGA
CGACTATCGGTATATGACGTTGAACGTAAGGCTCGCGAATGA
GCTTTCGAGCAACTTTGAGATGCAATATGAGTTGTCGTGGCA
GACTATGGACTTGGACGCTCGTGGCTACGACGGGCGCCAG
GCTGCAAAAGGGGACTATTGGAAATTGACGCTTGCGCCGAC
TTTCAAGGCACAAACGGGTGACTTCTTCATCCGACCAGAACT
CCGGTTGTTCGCAACTTACATGAACTGGTCACGCGACCTGG
ATGACTTTTCGGCAACGGATGATTTCGGGCAAAAGGGTTTCA
AGAGCGGTGGCGATTGGCAACTTGGCGTGCAGATGGAAACT
TGGTTTTAG

| | |
|---|---|
| *lacI* | ATGAAACCAGTAACGTTATACGATGTCGCAGAGTATGCCGGT<br>GTCTCTTATCAGACCGTTTCCCGCGTGGTGAACCAGGCCAGC<br>CACGTTTCTGCGAAAACGCGGGAAAAAGTGGAAGCGGCGAT<br>GGCGGAGCTGAATTACATTCCCAACCGCGTGGCACAACAACT<br>GGCGGGCAAACAGTCGTTGCTGATTGGCGTTGCCACCTCCAG<br>TCTGGCCCTGCACGCGCCGTCGCAAATTGTCGCGGCGATTAA<br>ATCTCGCGCCGATCAACTGGGTGCCAGCGTGGTGGTGTCGAT<br>GGTAGAACGAAGCGGCGTCGAAGCCTGTAAAGCGGCGGTGC<br>ACAATCTTCTCGCGCAACGCGTCAGTGGGCTGATCATTAACTA<br>TCCGCTGGATGACCAGGATGCCATTGCTGTGGAAGCTGCCTG<br>CACTAATGTTCCGGCGTTATTTCTTGATGTCTCTGACCAGACA<br>CCCATCAACAGTATTATTTTCTCCCATGAGGACGGTACGCGAC<br>TGGGCGTGGAGCATCTGGTCGCATTGGGTCACCAGCAAATCG<br>CGCTGTTAGCGGGCCCATTAAGTTCTGTCTCGGCGCGTCTGC<br>GTCTGGCTGGCTGGCATAAATATCTCACTCGCAATCAAATTCA<br>GCCGATAGCGGAACGGGAAGGCGACTGGAGTGCCATGTCCG<br>GTTTTCAACAAACCATGCAAATGCTGAATGAGGGCATCGTTCC<br>CACTGCGATGCTGGTTGCCAACGATCAGATGGCGCTGGGCG<br>CAATGCGCGCCATTACCGAGTCCGGGCTGCGCGTTGGTGCG<br>GATATCTCGGTAGTGGGATACGACGATACCGAAGATAGCTCA<br>TGTTATATCCCGCCGTTAACCACCATCAAACAGGATTTTCGCC<br>TGCTGGGGCAAACCAGCGTGGACCGCTTGCTGCAACTCTCTC<br>AGGGCCAGGCGGTGAAGGGCAATCAGCTGTTGCCAGTCTCA<br>CTGGTGAAAAGAAAAACCACCCTGGCGCCCAATACGCAAACC<br>GCCTCTCCCCGCGCGTTGGCCGATTCATTAATGCAGCTGGCA<br>CGACAGGTTTCCCGACTGGAAAGCGGGCAGTGA |
| *mScarlet-I* | ATGGTCAGTAAAGGAGAAGCTGTAATTAAAGAGTTTATGCGC<br>TTCAAAGTGCATATGGAAGGTTCCATGAACGGACATGAGTTC<br>GAGATTGAAGGTGAAGGTGAAGGACGCCCCTACGAGGGCAC<br>TCAAACTGCAAAGTTAAAGGTTACAAAAGGAGGACCTCTGCC<br>ATTTTCGTGGGACATTCTGAGCCCGCAGTTTATGTACGGCAG<br>CCGCGCGTTCATCAAACATCCCGCTGACATCCCAGACTATTA<br>CAAACAATCTTTCCCCGAAGGCTTTAAATGGGAACGCGTGAT<br>GAACTTTGAAGATGGTGGCGCTGTGACTGTGACCCAGGACA<br>CTTCATTAGAAGATGGAACCCTGATTTACAAGGTAAAGCTGC<br>GCGGCACCAACTTTCCCCCTGACGGACCTGTAATGCAGAAA<br>AAAACAATGGGTTGGGAGGCTAGTACAGAGCGTTTATACCCT<br>GAGGACGGTGTCTTAAAAGGAGACATCAAGATGGCGTTACG<br>TCTTAAGGATGGTGGTCGCTATTTAGCTGACTTCAAGACCAC<br>TTATAAAGCAAAGAAGCCCGTCCAAATGCCTGGAGCTTATAA |

| | | |
|---|---|---|
| | | CGTTGACCGTAAGTTAGACATCACCTCACATAACGAGGATTA<br>CACAGTTGTCGAACAGTATGAGCGCTCAGAAGGCCGTCATT<br>CGACTGGTGGAATGGACGAACTGTATAAATAA |
| | *sfgfp* | ATGCGTAAAGGCGAAGAGCTGTTCACTGGTGTCGTCCCTATT<br>CTGGTGGAACTGGATGGTGATGTCAACGGTCATAAGTTTTCC<br>GTGCGTGGCGAGGGTGAAGGTGACGCAACTAATGGTAAACT<br>GACGCTGAAGTTCATCTGTACTACTGGTAAACTGCCGGTACC<br>TTGGCCGACTCTGGTAACGACGCTGACTTATGGTGTTCAGTG<br>CTTTGCTCGTTATCCGGACCATATGAAGCAGCATGACTTCTT<br>CAAGTCCGCCATGCCGGAAGGCTATGTGCAGGAACGCACGA<br>TTTCCTTTAAGGATGACGGCACGTACAAAACGCGTGCGGAA<br>GTGAAATTTGAAGGCGATACCCTGGTAAACCGCATTGAGCTG<br>AAAGGCATTGACTTTAAAGAAGACGGCAATATCCTGGGCCAT<br>AAGCTGGAATACAATTTTAACAGCCACAATGTTTACATCACCG<br>CCGATAAACAAAAAAATGGCATTAAAGCGAATTTTAAAATTCG<br>CCACAACGTGGAGGATGGCAGCGTGCAGCTGGCTGATCACT<br>ACCAGCAAAACACTCCAATCGGTGATGGTCCTGTTCTGCTGC<br>CAGACAATCACTATCTGAGCACGCAAAGCGTTCTGTCTAAAG<br>ATCCGAACGAGAAACGCGATCATATGGTTCTGCTGGAGTTCG<br>TAACCGCAGCGGGCATCACGCATGGTATGGATGAACTGTAC<br>AAATGA |
| Plant Promoter | proM1 | ATAGAAAAGTTGGCTGCGGCCGCGAATTTCTGTCGATCTCTA<br>AATAATCGGGTCTAAGCTAGCTAGGCCACCCACAATCCCACT<br>ATCCTTTCGCAAGACCCTTCCTCTATATAAGGAAGTTCATTTC<br>ATTTGGAGAGAACACGGGGGACTCTAGACACC |
| | pro1x | ATAGAAAAGTTGGCTGCGGCCGCGAATTTCTATCGATCTATA<br>GATAATCGGGTCTAAGCTAGCTAGGCCACCCACAATCCCACT<br>ATCCTTTCGCAAGACCCTTCCTCTATATAAGGAAGTTCATTTC<br>ATTTGGAGAGAACACGGGGGACTCTAGACACC |
| | pro2x | ATAGAAAAGTTGGCTGCGGCCGCGAATTTCTATCGATCTATA<br>GATAATAGTTTCTATCGATCTATAGATAATCGGGTCTAAGCTA<br>GCTAGGCCACCCACAATCCCACTATCCTTTCGCAAGACCCTT<br>CCTCTATATAAGGAAGTTCATTTCATTTGGAGAGAACACGGG<br>GGACTCTAGACACC |
| | pro3x | ATAGAAAAGTTGGCTGCGGCCGCGAATTTCTATCGATCTATA<br>GATAATGTTTTCTATCGATCTATAGATAATAGTTTCTATCGATC<br>TATAGATAATCGGGTCTAAGCTAGCTAGGCCACCCACAATCC<br>CACTATCCTTTCGCAAGACCCTTCCTCTATATAAGGAAGTTCA<br>TTTCATTTGGAGAGAACACGGGGGACTCTAGACACC |
| | pro4x | ATAGAAAAGTTGGCTGCGGCCGCGAATTTCTATCGATCTATA<br>GATAATGTTTTCTATCGATCTATAGATAATGTTTTCTATCGATC<br>TATAGATAATAGTTTCTATCGATCTATAGATAATCGGGTCTAA<br>GCTAGCTAGGCCACCCACAATCCCACTATCCTTTCGCAAGAC<br>CCTTCCTCTATATAAGGAAGTTCATTTCATTTGGAGAGAACAC<br>GGGGGACTCTAGACACC |
| | pro5x | ATAGAAAAGTTGGCTGCGGCCGCGAATTTCTATCGATCTATA<br>GATAATGTTTTCTATCGATCTATAGATAATGTTTTCTATCGATC<br>TATAGATAATAGTTTCTATCGATCTATAGATAATAGTTTCTATC |

GATCTATAGATAATCGGGTCTAAGCTAGCTAGGCCACCCACA
ATCCCACTATCCTTTCGCAAGACCCTTCCTCTATATAAGGAA
GTTCATTTCATTTGGAGAGAACACGGGGGACTCTAGACACC

pro6x

ATAGAAAAGTTGGCTGCGGCCGCGAATTTCTATCGATCTATA
GATAATGATTTCTATCGATCTATAGATAATCATTTCTATCGATC
TATAGATAATCATTTCTATCGATCTATAGATAATGTTTTCTATC
GATCTATAGATAATAGTTTCTATCGATCTATAGATAATCGGGT
CTAAGCTAGCTAGGCCACCCACAATCCCACTATCCTTTCGCA
AGACCCTTCCTCTATATAAGGAAGTTCATTTCATTTGGAGAGA
ACACGGGGGACTCTAGACACC

proPIN2

ATCTTTCAATAGTTTCATCCTGTTTTATCAGGCTACATTCACTT
TGGTTGTTCTCTTTCAATTCATGAGTTAACAAAAAAACATGGA
TAAAGTTTAAAACTGAGATCACTTATTAAAGGCTTTTATTCTTC
TTCCTCTTGTGAAGATGTAAAGAAGAACAAACTCTTTTTAGGT
TTCTTGTTGGTCAATTCACCGTTTTTTTTAAAGAGATATAAGA
AATCGCGATGATCGTGTAGATGACATAGTGACCAACGAATTG
ATGGAGTCTTTGCTACGGATTGTGGAAAACCATCTCTTCGAT
ACATATTGGAACTTCAAAAGAGATTTTAGTGATAATCTAGTTA
GTATCTCCATCGTTAGATCCATAGGTCCATTAGGCCCAAACG
CCCAATCACGTGGCGTGATAAATACGTTAACAATTGCAATAT
GAAGAAAACACGACAATTAAATATGGAAGAGAGGCTAATTAT
TCATCGGTCCAATTGCCAATGTCTTTGACTGTAGTTAATAATA
AATCATTAAGAGATTTTCGTTTAAAAACAAAAATACACCGTTT
CAAGATTAAAAGATAAGTTAATTACAATTGTATTTTTCCTGATT
AAAAATCAGATGTTTACAGGGAACCGCAGATCGCTCAGAAAG
TGTAGTAAGATGCTCGCAAAAACCATGTGCATGTCCATCAAC
CTTCGCCGTCTCAATAGCTTACACAAAAATCGTTTTTGTTTTC
TTATTAATATAAAACCTAATCTAAATTTACAAATTTGTTTTCTTA
TTAATATAAAATCTAATCTAAATTTACAAATTTGTATTACACGT
ACAAGTGGATCCAAGTATCTGACTCTACAGATATTTTGAATCT
AACATTTGTGTAATTATAATTTATAATCTCTGAAGCATTTAATT
TTACTTACAAATAATGTTAGACCACGAGACAGAAGGTAAACC
AATCTTCCTTATCCCAAGATAAGATTCTTCAAATTAAATATAAC
GGCTTTTTGCAGAAGTAATAAAGATAGAAGGAATATCTAGTAT
CAACGGAAAAAAAGAAAAAATCTAGAGCACTTCGAAAAATTAT
GGACCAAAGAAGATTCAGACTGAAATTATTACTGTATTAGTAT
TTGTCCATTCACAAAATGCCGAGGAAGAAAAAAGGCATTTTG
GTTTATATTTTGTTTATTTGATATTCAAATGTCCAACGATCCTC
TCTAGCTAAGCTTAGCTATAATTCAATGTTTGAACACGAATCC
CATTATTTTAACACAAACAACATTAATTAAATATCGTCTCAAG
GAACTTCACTTCCTTGTCCATAAATACGTTATTTCACACCACA
TATACTCATCTATATCTCTATTTTTCTTCTTCTTCTCTCTCTCG
CCGGAAAAAGTAAATCAAACACC

proPYK10C.2

CTGCAACGAAGTGTACCAACAACTTGACTAGGATTCTAAGTT
CTTTTATGTATAGGATGTCTATATTAAACTACCATGACTAACA
TATATATAGTAGTTCCATATGCTCGATAAACTATGATAGATCA
ACAATTTTAAACATATAGTTTAACACTATTTATTTGTTCAACGT
CAATAGTTTATAGTTCGCATGCGCTCGGCTTAGATTTGGTCC
CCAACAGTCGAAATTGTCAAATAATATAAAATAAAAGTTTCAT
TGTTAGGATTCATTTATTCTTCGGGTGGTTATTGTAATAAAAG
GCAAAAGAAAAAGAAGAACAAAATTCACAAGTAAAAAAAAAG
ATAACATCATTCTTTTAGTCGACAAAAAAATAAAAAAAATCAAA

| | | |
|---|---|---|
| | | AAGATTTATTCAGTACTACAGTTTAATATTGTTTTGACTTTTTT<br>CTTTTTCTTTATATTATCTGAAAATTCTAGACTGCAGCTGAAA<br>CATGTGATATGGATTAAAGGCGTATCCAGTATCCACAGAAAG<br>AGGAGTGGTGTCGCTCACCCAGTCACCCTTGTTACTTGTTAG<br>ATAGCATTAATACATTTGTAAGCAACAGCTTATCTAATAGACA<br>TGTCTTAATTGGGAAATATGCTCTAAGATGATACAACCATGGT<br>TCCAACTGTTGACCACCATAACTGATAACATGTTGATTACATT<br>TTTTCTTTTCAGTTACAACGATTACTTTTTTGGGGAAATTATTG<br>ATATAATATGATTCATTGGATGATCCGATATCATGCATATAAA<br>GTTGTATCTCGTGAAACACGAGATAGTATTATACTCCATTCTT<br>TCATTATCGGAGTATGTTTAAAATTTGAAAACAAATACAGACA<br>CGGACCGTGGTCTTTACCTTCAGAAAAAAAAAGAGAAAAAAA<br>AACAATCCACTGTTTATTATAGGAGTTGTAGAAAATCGGGCA<br>ACGATATTCGATATGAGTTATTATTAGGGCCTTATTATTATAT<br>GGTATTACTGGATATTACTAAATAATCATATAAATATCACATTT<br>TAATATACACTCGTTGGACACGCGGAATATTATATGTTCTAAA<br>TGTTAAAAAATCAACAGAATACAACGATCGACGGATCTAGAG<br>TCTAGACCATGCAAATACCTCATCCTATTTACATATAATAACT<br>GTGCATATAGTTTAGTCAAATAAAAAGGTAAAGAAACAATATA<br>CAACCTATAACGTCAATATCCATGTACGTAGTAATAATTAGGA<br>TATGACACGAACACACGATATCTTGATATATACAAAATGAAAA<br>CTTAAAAATTGATTAATATGGCCTGGCTGGGTATATTATTAAA<br>AAAACATAAAGAGAGATCAATAATTGATTCGAAGATCACTATA<br>TAAAGAACGTCTACGATATGTAAAAGAACCATCCTAAACATTT<br>TTTCTTGAATAAAATCAGAATTACAAACAAAACACC |
| Plant Gene | *SWEET11-mTurquoise2* | ATGGGTTCAATGAGTCTCTTCAACACTGAAAACACATGGGCC<br>TTTGTCTTTGGCTTGCTCGGCAACCTTATCTCCTTTGCCGTGT<br>TCCTATCTCCTGTGCCAACGTTCTATAGGATTTGGAAGAAAA<br>AGACAACAGAAGGGTTTCAGTCTATTCCTTATGTTGTGGCGC<br>TCTTCAGTGCGACGCTTTGGCTTTACTATGCGACACAGAAGA<br>AAGATGTATTCCTCCTCGTAACCATTAACGCCTTTGGTTGCTT<br>CATCGAAACCATCTACATCTCTATGTTCCTTGCCTACGCTCC<br>CAAGCCAGCTCGGATGTTGACAGTGAAGATGCTACTTCTTAT<br>GAACTTTGGAGGATTCTGTGCGATTCTCCTTCTTTGCCAATTC<br>TTGGTAAAAGGAGCCACACGTGCTAAGATTATCGGAGGAATC<br>TGTGTCGGATTCTCTGTTTGTGTTTTCGCTGCTCCTCTAAGCA<br>TAATCAGGACGGTAATAAAGACAAGAAGTGTGGAGTACATGC<br>CCTTTAGCTTATCCTTAACCCTTACCATCAGTGCTGTCATATG<br>GCTCCTTTATGGTCTTGCTCTCAAGGACATCTATGTTGCTTTC<br>CCGAATGTGCTTGGTTTTGCTCTCGGTGCACTCCAAATGATA<br>CTCTACGTTGTCTACAAATACTGTAAAACGTCGCCGCATCTA<br>GGAGAGAAAGAAGTCGAAGCTGCTAAGTTACCGGAGGTGAG<br>CCTCGATATGTTGAAGCTAGGCACAGTTTCATCCCCTGAGCC<br>AATCTCAGTGGTTCGTCAAGCGAACAAGTGTACCTGCGGAAA<br>TGATCGAAGGGCTGAGATTGAAGATGGACAAACCCCTAAACA<br>TGGCAAGCAGTCCTCTTCCGCAGCAGCTACAGGAGGAGGGA<br>TGGTGAGCAAGGGAGAAGAACTGTTTACCGGAGTTGTTCCC<br>ATCCTGGTTGAGCTGGACGGTGACGTAAACGGACACAAGTT<br>CAGCGTTTCCGGAGAGGGAGAAGGAGATGCAACCTACGGCA<br>AGCTGACCCTGAAGTTCATCTGCACCACTGGCAAGCTGCCC<br>GTTCCCTGGCCCACCCTCGTTACTACCCTGTCTTGGGGTGT<br>GCAGTGCTTCGCCCGCTACCCCGACCACATGAAGCAGCACG<br>ACTTCTTCAAGTCCGCCATGCCCGAAGGCTACGTCCAGGAG<br>CGCACCATCTTCTTCAAGGACGACGGCAACTACAAGACCCG |

| | |
|---|---|
| | CGCCGAGGTGAAGTTCGAGGGCGACACCCTGGTGAACCGC<br>ATCGAGCTGAAGGGCATCGACTTCAAGGAGGACGGCAACAT<br>CCTGGGGCACAAGCTGGAGTACAACTACTTTAGCGACAACG<br>TCTATATCACCGCCGACAAGCAGAAGAACGGCATCAAGGCC<br>AACTTCAAGATCCGCCACAACATCGAGGACGGTGGAGTTCA<br>GCTTGCTGATCACTACCAACAGAACACCCCTATCGGTGACG<br>GACCAGTTCTGCTTCCAGATAACCATTACCTTTCTACCCAGT<br>CCAAGCTGAGCAAAGACCCCAACGAGAAGCGCGATCACATG<br>GTCCTGCTGGAGTTCGTGACCGCCGCCGGGATCACTCTCGG<br>CATGGACGAGCTGTACAAGTAA |
| *SWEET11-mTurquoise2 intron* | ATGGGTTCAATGAGTCTCTTCAACACTGAAAACACATGGGCC<br>TTTGTCTTTGGCTTGCTCGGCAACCTTATCTCCTTTGCCGTGT<br>TCCTATCTCCTGTGCCAACGTTCTATAGGATTTGGAAGAAAA<br>AGACAACAGAAGGGTTTCAGTCTATTCCTTATGTTGTGGCGC<br>TCTTCAGTGCGACGCTTTGGCTTTACTATGCGACACAGAAGA<br>AAGATGTATTCCTCCTCGTAACCATTAACGCCTTTGGTTGCTT<br>CATCGAAACCATCTACATCTCTATGTTCCTTGCCTACGCTCC<br>CAAGCCAGCTCGGATGTTGACAGTGAAGATGCTACTTCTTAT<br>GAACTTTGGAGGATTCTGTGCGATTCTCCTTCTTTGCCAATTC<br>TTGGTAAAAGGAGCCACACGTGCTAAGATTATCGGAGGAATC<br>TGTGTCGGATTCTCTGTTTGTGTTTTCGCTGCTCCTCTAAGCA<br>TAATCAGGACGGTAATAAAGACAAGAAGTGTGGAGTACATGC<br>CCTTTAGCTTATCCTTAACCCTTACCATCAGTGCTGTCATATG<br>GCTCCTTTATGGTCTTGCTCTCAAGGACATCTATGTTGCTTTC<br>CCGAATGTGCTTGGTTTTGCTCTCGGTGCACTCCAAATGATA<br>CTCTACGTTGTCTACAAATACTGTAAAACGTCGCCGCATCTA<br>GGAGAGAAAGAAGTCGAAGCTGCTAAGTTACCGGAGGTGAG<br>CCTCGATATGTTGAAGCTAGGCACAGTTTCATCCCCTGAGCC<br>AATCTCAGTGGTTCGTCAAGCGAACAAGTGTACCTGCGGAAA<br>TGATCGAAGGGCTGAGATTGAAGATGGACAAACCCCTAAACA<br>TGGCAAGCAGTCCTCTTCCGCAGCAGCTACAGGAGGAGGGA<br>TGGTGAGCAAGGGAGAAGAACTGTTTACCGGAGTTGTTCCC<br>ATCCTGGTTGAGCTGGACGGTGACGTAAACGGACACAAGTT<br>CAGCGTTTCCGGAGAGGGAGAAGGAGATGCAACCTACGTAA<br>GTTTCTGCTTCTACCTTTGATATATATATAATAATTATCATTAA<br>TTAGTAGTAATATAATATTTCAAATATTTTTTTCAAAATAAAAG<br>AATGTAGTATATAGCAATTGCTTTTCTGTAGTTTATAAGTGTG<br>TATATTTTAATTTATAACTTTTCTAATATATGACCAAAATTTGTT<br>GATGTGCAGGGCAAGCTGACCCTGAAGTTCATCTGCACCAC<br>TGGCAAGCTGCCCGTTCCCTGGCCCACCCTCGTTACTACCC<br>TGTCTTGGGGTGTGCAGTGCTTCGCCCGCTACCCCGACCAC<br>ATGAAGCAGCACGACTTCTTCAAGTCCGCCATGCCCGAAGG<br>CTACGTCCAGGAGCGCACCATCTTCTTCAAGGACGACGGCA<br>ACTACAAGACCCGCGCCGAGGTGAAGTTCGAGGGCGACACC<br>CTGGTGAACCGCATCGAGCTGAAGGGCATCGACTTCAAGGA<br>GGACGGCAACATCCTGGGGCACAAGCTGGAGTACAACTACT<br>TTAGCGACAACGTCTATATCACCGCCGACAAGCAGAAGAAC<br>GGCATCAAGGCCAACTTCAAGATCCGCCACAACATCGAGGA<br>CGGTGGAGTTCAGCTTGCTGATCACTACCAACAGAACACCC<br>CTATCGGTGACGGACCAGTTCTGCTTCCAGATAACCATTACC<br>TTTCTACCCAGTCCAAGCTGAGCAAAGACCCCAACGAGAAG<br>CGCGATCACATGGTCCTGCTGGAGTTCGTGACCGCCGCCGG<br>GATCACTCTCGGCATGGACGAGCTGTACAAGTAA |

| | |
|---|---|
| *mTurquoise2* | ATGGGGATGGTGAGCAAGGGAGAAGAACTGTTTACCGGAGT<br>TGTTCCCATCCTGGTTGAGCTGGACGGTGACGTAAACGGAC<br>ACAAGTTCAGCGTTTCCGGAGAGGGAGAAGGAGATGCAACC<br>TACGGCAAGCTGACCCTGAAGTTCATCTGCACCACTGGCAA<br>GCTGCCCGTTCCCTGGCCCACCCTCGTTACTACCCTGTCTTG<br>GGGTGTGCAGTGCTTCGCCCGCTACCCCGACCACATGAAGC<br>AGCACGACTTCTTCAAGTCCGCCATGCCCGAAGGCTACGTC<br>CAGGAGCGCACCATCTTCTTCAAGGACGACGGCAACTACAA<br>GACCCGCGCCGAGGTGAAGTTCGAGGGCGACACCCTGGTG<br>AACCGCATCGAGCTGAAGGGCATCGACTTCAAGGAGGACGG<br>CAACATCCTGGGGCACAAGCTGGAGTACAACTACTTTAGCGA<br>CAACGTCTATATCACCGCCGACAAGCAGAAGAACGGCATCA<br>AGGCCAACTTCAAGATCCGCCACAACATCGAGGACGGTGGA<br>GTTCAGCTTGCTGATCACTACCAACAGAACACCCCTATCGGT<br>GACGGACCAGTTCTGCTTCCAGATAACCATTACCTTTCTACC<br>CAGTCCAAGCTGAGCAAAGACCCCAACGAGAAGCGCGATCA<br>CATGGTCCTGCTGGAGTTCGTGACCGCCGCCGGGATCACTC<br>TCGGCATGGACGAGCTGTACAAGTAA |
| *AmtR-ERF2-NLS* | ATGGCTGGCGCCGTGGGCAGACCCAGAAGATCTGCTCCTCG<br>GAGAGCCGGCAAGAACCCCCGGGAAGAGATTCTGGATGCCA<br>GCGCCGAGCTGTTCACCAGACAGGGCTTTGCCACCACCAGC<br>ACCCACCAGATTGCCGACGCTGTGGGCATCAGACAGGCCAG<br>CCTGTACTACCACTTCCCCAGCAAGACCGAGATCTTCCTGAC<br>CCTGCTGAAAAGCACCGTGGAACCCTCCACCGTGCTGGCCG<br>AGGATCTGTCTACCCTGGACGCCGGACCCGAAATGAGACTG<br>TGGGCTATCGTGGCCAGCGAAGTGCGGCTGCTGCTGAGCAC<br>CAAGTGGAACGTGGGCCGGCTGTACCAGCTGCCCATCGTGG<br>GCTCTGAGGAATTCGCCGAGTACCACAGCCAGCGCGAGGCC<br>CTGACCAACGTGTTCAGAGATCTGGCCACCGAGATTGTGGG<br>CGACGACCCCAGAGCCGAACTGCCCTTCCACATCACCATGA<br>GCGTGATCGAGATGCGGCGGAACGACGGCAAGATCCCTAG<br>CCCTCTGAGCGCCGACTCTCTGCCCGAGACAGCCATTATGC<br>TGGCTGACGCCTCCCTGGCTGTGCTGGGAGCACCTCTGCCT<br>GCCGACAGAGTGGAAAAGACACTGGAACTGATCAAGCAGGC<br>CGACGCCAAGGGAGGAGGTGAATCCGACTACGCTTTGTTGG<br>AGTCGATAACACGTCACTTGCTAGGAGGAGGAGGAGAGAAC<br>GAGCTGCGACTCAATGAGTCAACACCGAGTTCGTGTTTCACA<br>GAGAGTTGGGGAGGTTTGCCATTGAAAGAGAATGATTCAGA<br>GGACATGTTGGTGTACGGACTCCTCAAAGATGCCTTCCATTT<br>TGACACGTCATCATCGGACTTGAGCTGTCTTTTTGATTTTCCG<br>GCGCCGAAGAAGAAGAGGAAGGTTTAG |

**Table S2.** Biological resources including bacterial strains/plasmids, plant material/seeds, and key DNA. Operator DNA sequences bound by biosensor repressor transcription factors are underlined in relevant bacterial promoters.
